# A Biologically Inspired Attention Model for Neural Signal Analysis

**DOI:** 10.1101/2024.08.13.607787

**Authors:** Nicolas Skatchkovsky, Natalia Glazman, Alexander Egea-Weiss, Sadra Sadeh, Florencia Iacaruso

**Author notes:** These authors contributed equally to this work (Sadra Sadeh, Florencia Iacaruso).

## Abstract

Understanding how the brain represents sensory information and triggers behavioural responses is a fundamental goal in neuroscience. Recent advances in neuronal recording techniques aim to progress towards this milestone, yet the resulting high-dimensional responses are challenging to interpret and link to relevant variables. Although existing machine learning models propose to do so, they often sacrifice interpretability for predictive power, effectively operating as black boxes. In this work, we introduce SPARKS, a biologically inspired model capable of high decoding accuracy and interpretable discovery within a single framework. SPARKS adapts the *self-attention* mechanism of large language models to extract information from the timing of single spikes and the sequence in which neurons fire using Hebbian learning. Trained with a criterion inspired by predictive coding to enforce temporal coherence, our model produces low-dimensional latent embeddings that are robust across sessions and animals. By directly capturing the underlying data distribution through a generative encoding-decoding framework, SPARKS matches or exceeds state-of-the-art decoders across diverse electrophysiology and calcium imaging datasets from the motor, visual and entorhinal cortices. Crucially, the Hebbian coefficients learned by the model are interpretable, allowing us to infer directed functional interactions and recover the known functional hierarchy of the mouse visual cortex. Overall, SPARKS unifies representation learning, high-performance decoding and model interpretability in a single framework by bridging neuroscience and AI, providing a powerful and versatile tool for dissecting neural computations and marking a step towards the next generation of biologically inspired intelligent systems.

---

Recent advancements in imaging and electrophysiology techniques have enabled neuroscientists to capture neuronal activity at an unprecedented scale^1^, yielding recordings comprising hundreds to thousands, and even a million neurons over extended periods^2–5^. Interpreting this neural activity and relating it to sensory, motor, and cognitive processes is a key goal of neuroscience^6^, yet this task remains challenging for a number of reasons. Neural recordings are high dimensional, both spatially – because of the number of neurons recorded, and temporally – because of high temporal resolution or long recording times, making them hard to interpret. In addition, a substantial portion of the variance can typically be attributed to behavioural parameters like uninstructed movements^7^, introducing a substantial amount of variability across trials, sessions and animals^8,9^ and masking the relationship between neuronal activity and phenomena of interest. This variability can also result from separate neuronal populations being recorded in different sessions, from changes in neural activity due to different brain states, or from effects such as neural drift^10^. Developing tools to tackle these challenges is now imperative to the field.

Finding suitable low-dimensional representations is a promising avenue to understand complex, high-dimensional neuronal data, as previous research indicates that most of the variance of the neural activity typically lies within low-dimensional subspaces known as *neural manifolds*^11–13^, and a growing body of evidence suggests that the prism of neural manifolds provides information that cannot be found at the level of single neurons^14,15^. Early studies of population dynamics relied on techniques such as principal components analysis (PCA), which ignores the temporal nature of neural processes and is typically applied to binned or averaged data, leading to discard fine-grained information^11,12^. Subsequent machine learning (ML) approaches demonstrated the benefits of variational autoencoders (VAEs) to generate low-dimensional representations of the data and perform prediction^16^, and modern techniques often rely on deep learning models such as Transformers, sacrificing interpretability for predictive power^17–22^. In parallel, contrastive learning has become a successful approach to perform representation learning with temporal data^17^, although this approach ignores the specific sequential, asynchronous nature of events in neuronal recordings. Beyond the loss of crucial temporal information, many of these techniques also require extensive pre-processing and manual data curation, limiting standardisation and reproducibility^23^, and few are capable of finding common representations of the data recorded across sessions and animals^17^. In this work, we propose to combine the generative, exploratory capabilities of VAEs^24,25^, and the high predictive power of Transformers to improve our understanding of neural data.

Two properties of neural recordings are poorly captured by existing sequence models. First, information is carried by the relative timing and order of sparse, asynchronous spikes across pairs of neurons, whereas the dot-product attention of Transformers compares tokens presented simultaneously and relies on positional embeddings to recover their order. Second, neural and behavioural states evolve smoothly, so useful representations should predict future activity rather than only describe the present. Motivated by the growing literature on physics- and biology-inspired AI models, which promises to develop more powerful and explainable AI^26^, our contributions tackle these limitations in two ways. Firstly, we integrate Transformers into VAEs and adapt them to the unique nature of neural data using one of the processes behind learning in biological brains. By designing a *Hebbian attention layer* to operate directly on sparse populations of asynchronous events, we developed a computational framework that is better matched to the structure of neural data than the components of a standard Transformer. Unlike black-box deep learning models, the coefficients learned by this model are reflective of functional interactions between neurons, allowing us to gain biological insights about network interactions. Secondly, we introduce a novel criterion inspired by predictive coding, which maximises the probability for the model to predict the correct output for a number of time-steps in the future, given the past latent embeddings. Using this combination, we obtain state-of-the-art predictive power and robust latent representations to analyse neural recordings. Latent embeddings obtained from recordings in different sessions, animals or recording sites can be directly compared within the same latent space, which will be key towards the effort of building a brain-wide understanding of neural mechanisms.

The proposed model is a Sequential Predictive Autoencoder for the Representation of spiKing Signals (SPARKS). Our approach offers several key advantages: training directly on spiking signals or calcium imaging data with minimal preprocessing; dynamically faithful low-dimensional latent embeddings; state-of-the-art predictive power; unsupervised exploration of the data; coefficients in the Hebbian attention layer that can be linked to functional relationships between neurons; and robust encodings across repetitions, sessions, animals, and brain areas. Throughout, we showcase SPARKS’ capabilities using diverse datasets, highlighting its potential to uncover biological results.

## SPARKS: A Variational Autoencoder with Hebbian Attention

To obtain consistent latent embeddings that capture most of the variance from high dimensional neuronal responses, SPARKS integrates into the encoder of a VAE a Transformer architecture implementing our Hebbian attention layer (Figure 1A-C). Following the seminal work^27^, the encoder comprises several attention blocks, each composed of an attention layer, followed by a feedforward network (Figure 1C, Figure S1A). The first attention block implements the Hebbian self-attention mechanism that we detail in the next section, with the following blocks implementing conventional dot-product attention with rotary embeddings^28^ (Figure S1A). To obtain latent representations of the input data, the output of the last attention block is projected into the desired latent dimension. Throughout, the decoder is a linear layer or multilayer perceptron (MLP), tasked to predict a desired *reference* signal. The reference signal can be chosen as a general supervisory signal such as behavioural data or sensory inputs to perform supervised learning, or as the input spike train itself to perform unsupervised learning. As we will demonstrate, both approaches can be used in complement of each other. Training via unsupervised learning allows the exploration of low dimensional representations of the data produced by the encoder without imposing on the model prior information about task variables. This is particularly interesting when investigating how complex combinations of external and internal parameters are encoded in the data. This analysis can be complemented by supervised training to understand how external parameters recorded during the experiments are encoded in the neural activity. In the supervised mode, the latent space is shaped by the reference signal by design, isolating the part of the neural manifold that relates to the variable of interest, whereas unsupervised training captures the manifold as a whole.

**Figure 1.**
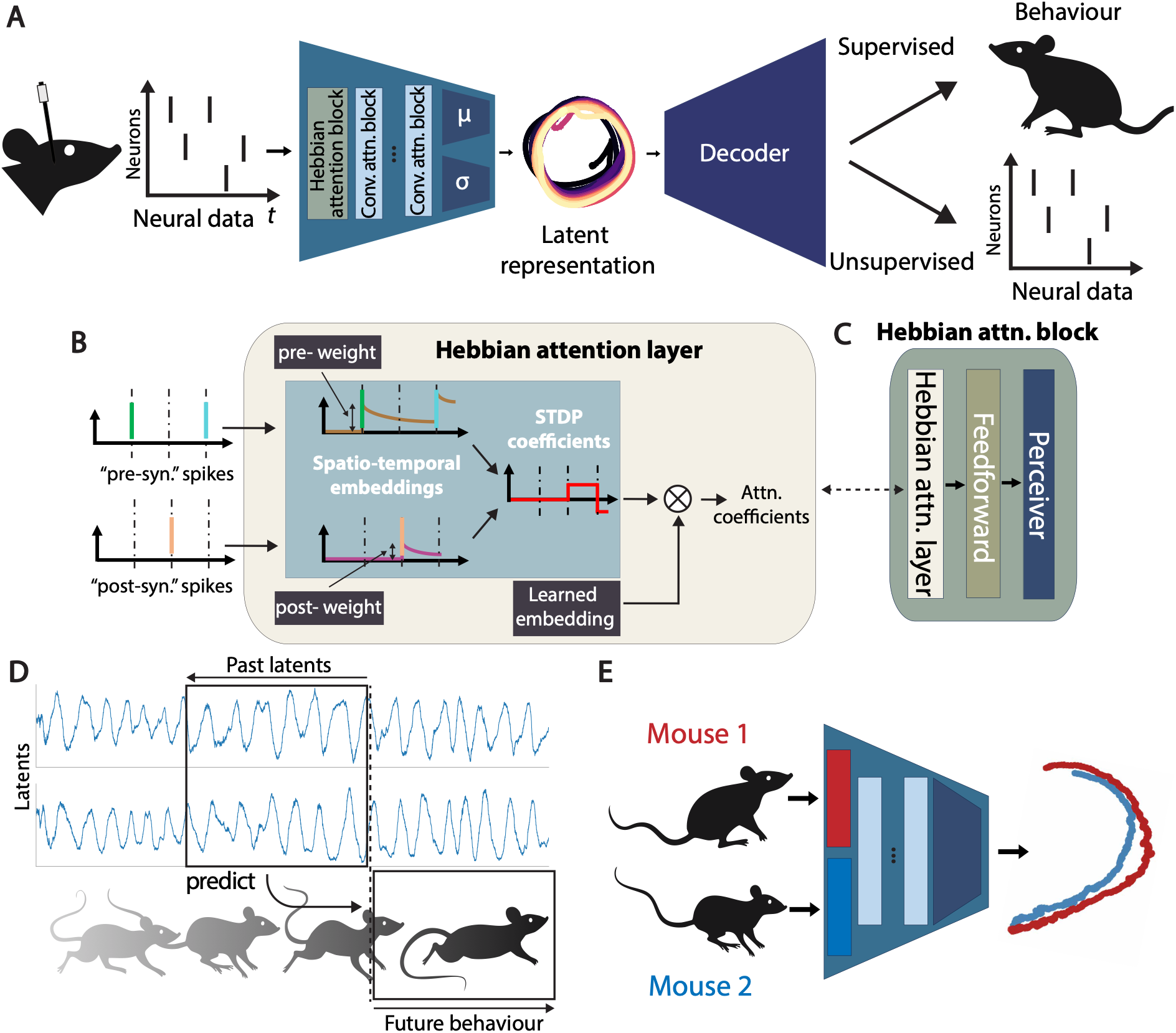
Overview of SPARKS. (**A**) Architecture of the proposed variational autoencoder operating directly on spiking signals. Blocks **µ**(⋅) and **σ**(⋅) represent the mean and standard deviation of the latent distribution. (**B**) Detailed view of the Hebbian attention layer. (**C**) Schematic of the Hebbian attention block comprising an Hebbian attention layer, feedforward network, and spatial perceiver block. (**D**) Illustration of the predictive conditional distribution: at every time-step, a window τ_f_ of past latent embeddings is used to predict the future τ_p_ samples from the reference signal. (**E**) Illustration of training with data from multiple sessions or animals. SPARKS learns a different Hebbian attention layer for each session, while keeping the rest of the encoding network shared.

We train SPARKS through a novel objective which is obtained by extending the definition of causally conditioned distributions to the framework of predictive coding^29,30^. Causally conditioned distributions model the probability of a random variable (RV) given its own past and that of other RVs of interest: in this context, we maximise the probability to output the desired reference signal given the latent embedding produced by the encoder. More specifically, we introduce the *predictive causally conditioned (PCC)* distribution, which consists in maximising the likelihood of future samples from the reference signal given a window of past samples from the latent space (Figure 1D). The use of the PCC is inspired by the predictive coding theory, namely, that the encodings produced by the brain’s internal generative model constantly aim to predicting future sensory inputs^31^. By forcing the model to learn representations that are predictive of longer-term future dynamics, the proposed objective introduces temporal coherence in the representations produced to capture the true underlying generative process of the data. This aligns with recent discoveries in Transformers, where training models to predict multiple future tokens simultaneously has been shown to improve both performance and training efficiency^32^, and we find that conditioning predictions on a window of past latent embeddings, together with a longer prediction horizon, improves temporal models of neural data.

### Applying Self-Attention to Neural Data

Transformers are powered by *self-attention* mechanisms^27^, which operate in two steps. Firstly, they learn positional encodings to make sense of the order of *tokens* in the sequence, before computing relationships between these tokens in order for the model to determine which are relevant to one another. We propose that Hebbian mechanisms offer an implementation of attention for neural sequences by fusing these two steps via the computation of synaptic strength coefficients based on the relative spike times of pre- and post-synaptic neurons. This *Hebbian attention* is a fundamentally spatiotemporal operator: it is sensitive to both which of the *N* neurons are firing (the spatial component) and when they are firing relative to one another (the temporal component, captured by dynamically evolving eligibility traces, Figure 1B). The Hebbian attention layer implements a recurrent process, which can be embedded into a VAE architecture to benefit from the previously demonstrated capabilities of recurrent VAEs for neural data analysis^16^. This is unlike conventional dot-product attention, which relies on various positional embedding techniques to introduce temporality across tokens presented simultaneously to the model^27^.

With Hebbian attention, we sequentially compute coefficients between neurons at every time instant via spike-time dependent plasticity (STDP), regardless of prior knowledge of connectivity. This computation is carried out by maintaining *eligibility traces* for each pair of neurons, which are calculated using weighted averages of the neurons’ spiking history: they are potentiated when either neuron spikes, and decay exponentially otherwise (Figure 1B). The time constants of decay and the update magnitudes for pre- and post-synaptic traces are initialised to values informed by experimental recordings^33^, and these parameters can optionally be learned for each neuron pair via backpropagation (SPARKS-learned). Although this configuration increases the computational cost of training our model, we will show it presents advantages for complex datasets, as compared to keeping these parameters fixed (SPARKS-fixed). We benchmarked the runtime and memory requirements of both configurations across a range of parameters (Methods, Figure S1D).

The coefficients obtained via STDP can be likened to the dot-product between keys and queries in the conventional dot-product attention^27^, and these values are then projected through a *D*-dimensional learned embedding (*D* ≪ *N*). Optionally, the model can implement several Hebbian heads with distinct decay time constants to capture correlations at different timescales. Following a feed-forward network with residual connections (Figure S1B), neuron embeddings are summed with a learned per-neuron embedding and compressed by a Perceiver-style module into *M*-dimensional representations (with *M* ≪ *N*, Methods, Figure S1C). This read-out decouples the dimensionality of the representation from the number of recorded neurons, and our model can be fitted jointly to several recordings with different neuron counts and identities (Figure 1E), with session-specific Hebbian layer and identity embeddings while all other parameters are shared. The resulting *M* × *D*-dimensional values are subsequently flattened and passed to conventional attention blocks. Our results focus on electrophysiology and deconvolved calcium data, but our method can also be adapted to calcium traces obtained through microscopy (Methods).

As we will demonstrate, the implementation of an attention mechanism via causal STDP enables the model to capture sequential information. It is well suited to encoding neural data where information lies in the precise order of spikes, such as in the entorhinal-hippocampal system^34^, as highlighted in the following section.

### Hebbian Attention Captures Sequential Information

We start by illustrating the capacity of our model to extract known low-dimensional latent structures using recordings from the medial entorhinal cortex^35^ (MEC, Figure 2A). It was recently demonstrated that oscillatory sequences are propagated in this brain region on the scale of tens of seconds to minutes in passive mice, and we recovered the circular latent topology found in a previous study by training SPARKS directly on the deconvoluted recordings (Figure 2B). From the latent trajectory, we retrieved the phase of the oscillations (Figure S2A-B, Methods), which allowed us to order neurons according to their correlation to the phase signal (Figure 2C). We found that sorting units based on SPARKS’ results improved upon previous work, as measured by the height of the peak in the power spectral density (PSD) after population decoding of the spike train (Figure 2C, Figure S2C-D, Methods). The signal recovered after sorting with SPARKS had a peak power of 7.4 *mV*^2^. *Hz*^−1^, outperforming previous methods relying on smooth firing rates^35^ (4.8 *mV*^2^. *Hz*^−1^).

**Figure 2.**
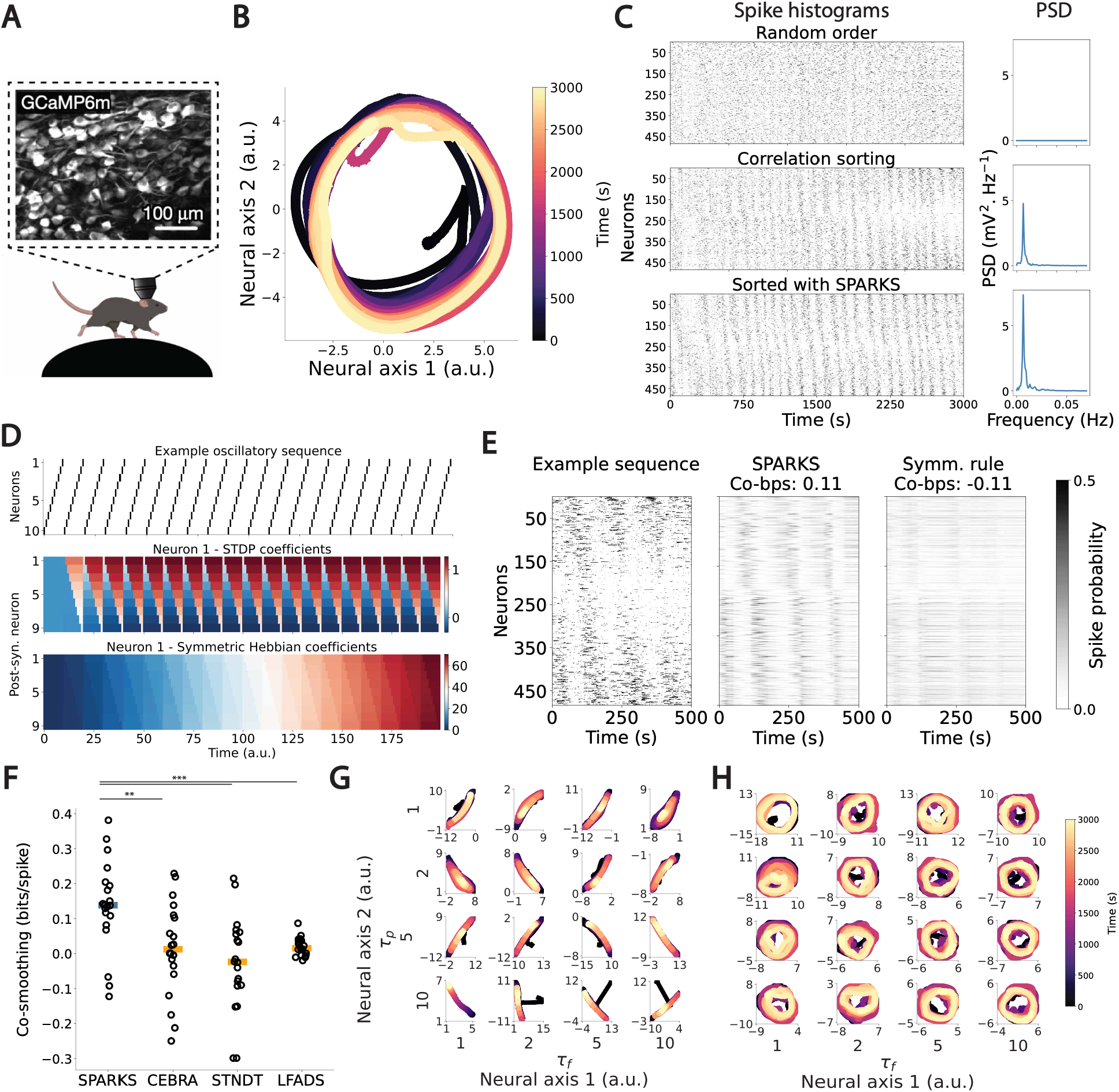
SPARKS reveals oscillatory sequences from MEC data. (**A**) Illustration of calcium recording in the MEC of head-fixed mice freely running on a wheel^35^. (**B**) Two-dimensional embeddings obtained from SPARKS-fixed for the whole session after unsupervised training. (**C**) Raster plot of thresholded calcium signals of all cells in an example session (bin size = 129 ms, n = 464 cells). Top: random order. Middle: Cells were sorted according to their correlation values with one arbitrary cell, in a descending manner. Bottom: Cells were sorted according to the peak of their correlation with the signal phase recovered with SPARKS-fixed. (**D**) Raster plot of synthetic firing patterns (top) and corresponding coefficients obtained after application of STDP (middle), and an alternative symmetric rule (bottom) (**E**) Raster plot for left-out data (left) and corresponding predicted spike probabilities after training with SPARKS-fixed (centre), and the benchmark symmetric rule (right). (**F**) Co-smoothing scores for SPARKS-fixed and competing models. Coloured lines represent the median over 20 runs, and each dot an individual run. (**G**) Evolution of the two-dimensional embeddings obtained from the model with the symmetric Hebbian rule for different values of τ_f_ and τ_p_. (**H**) Evolution of the two-dimensional embeddings obtained from SPARKS-fixed for different values of τ_f_ and τ_p_. *: p < 0.05, **: p < 0.01, ***: p < 0.001.

To motivate the choice of Hebbian attention and examine how its specific mode of operation enables SPARKS’ performance, we replaced the Hebbian attention layer with various alternative modules while leaving the downstream architecture unchanged. Firstly, we investigated an alternative, symmetric Hebbian rule in which the coefficients are potentiated regardless of the order of firing of the pre- and post-synaptic neurons (Methods). As illustrated on a synthetic oscillatory sequence with 10 neurons, applying STDP results in periodic coefficients, while the coefficients calculated with the symmetric rule diverged over time (Figure 2D). On the MEC data, the model implementing the symmetric rule could not predict the behaviour of neurons left out of its training data, as measured by a co-smoothing score calculating how well each model’s predictions capture fluctuations in the spiking sequence, compared to a model predicting average spike rates^36^ (Methods, Figure 2E, Figure S2E-F).

Although the symmetric Hebbian rule is inefficient in this case, it is plausible that keeping track of the neurons’ activity through an eligibility trace and processing the resulting values using a deep learning module might be sufficient to capture the simple dynamics of this data. We found it not to be the case: replacing the Hebbian block with a multilayer perceptron (MLP), a recurrent neural network (RNN) or a conventional attention layer lowered median co-smoothing, consistent with the idea that tracking the co-fluctuation between each pair of neurons contributes to the performance of Hebbian attention, although not all these differences reached significance (*p* = 0.02 − 0.12, two-sided Mann–Whitney U test; Figure S2F). Tracking co-fluctuation could still be implemented in a number of ways, and we replaced Hebbian attention with an alternative module measuring correlations between each pair of neurons over time (Methods). Interestingly, this model performed poorly, showing that Hebbian attention’s advantage lies in its capability to capture the precise spiking order across pairs of neurons, at a single-spike resolution. SPARKS also performed favourably against a range of competing deep learning models^16,17,22^ (Figure 2F).

Next, we evaluated the impact of the proposed learning rule. Increasing the value of the prediction window *τ*_*f*_ in the PCC allowed us to recover the circular structure of the latent space with the symmetric rule (Figure 2G) and increasing both *τ*_*f*_ and *τ*_*p*_ improved the corresponding model’s co-smoothing performance (Figure 2H), hinting at the PCC’s relevance to capture temporal dynamics. While we demonstrate in the following how the PCC is beneficial in the case of more complex dynamics, our model’s performance on this task was consistent across values of *τ*_*f*_ and *τ*_*p*_, and we found that choosing *τ*_*f*_ = *τ*_*p*_ = 1 was sufficient for SPARKS to capture the circular structure of the population responses. Increasing the past and future prediction windows was detrimental to the other modules with which we replaced Hebbian attention (Figure S2H). This result could be explained by increased parameter counts when varying *τ*_*f*_ and *τ*_*p*_. All modules having similar numbers of parameter to SPARKS, this suggests our method’s robustness to overfitting (Figure S2H-I). SPARKS-fixed was the best performing version of our model on this task, where a single trial was likely not sufficient to fit the learned configuration’s larger number of parameters (Figure S2H-I).

STDP can therefore be used as a directed, spike-time resolution spatio-temporal operator to capture fine-grained sequential information and recover the structure of neuronal populations. By conditioning predictions on a window of past latents, the PCC helps capture temporal structure when the encoder alone does not, as in the case of the symmetric rule.

### Dynamically Faithful, Low-Dimensional and Robust Encodings

To investigate how latent embeddings can be related to parameters of interest such as behavioural variables, we considered electrophysiology recordings in Brodmann’s area 2 of the somatosensory cortex in monkeys performing a centre-out task^37,38^ (Figure 3A). Training SPARKS on a single recording from the Area2 Bump dataset to predict the position of the animal’s hand, we found that the structure of the latent embeddings obtained closely matched the actual hand trajectories (Figure 3B). When training the model via unsupervised learning, despite not being informed by the animal’s behaviour, latent trajectories were still clearly separated across different trial conditions and matched the organisation in space of the target directions. Replacing the Hebbian attention layer with an alternative attention mechanism for neural data^20^ or the modules introduced in the previous section still produced latent embeddings matching the structure of the animal’s movements (Methods, Figure 3B, Figure S3A). In contrast, while the embeddings produced by the state-of-the-art model for manifold learning^17^ are clearly separated by trial direction, they do not visually correspond to the trajectories of the animal’s hand (Figure 3B). Quantitatively, we measured Procrustes^39^ distances to obtain a geometric measure of similarity between latent representations and the reference signal. Distances were lowest for SPARKS but only slightly higher for the POYO+, MLP, GRU and symmetric controls, mirroring their visual resemblance to the hand trajectories (Figure S3B). These results suggest that combining Transformer architectures with VAEs is a promising avenue to capture low-dimensional structure in neural data.

**Figure 3.**
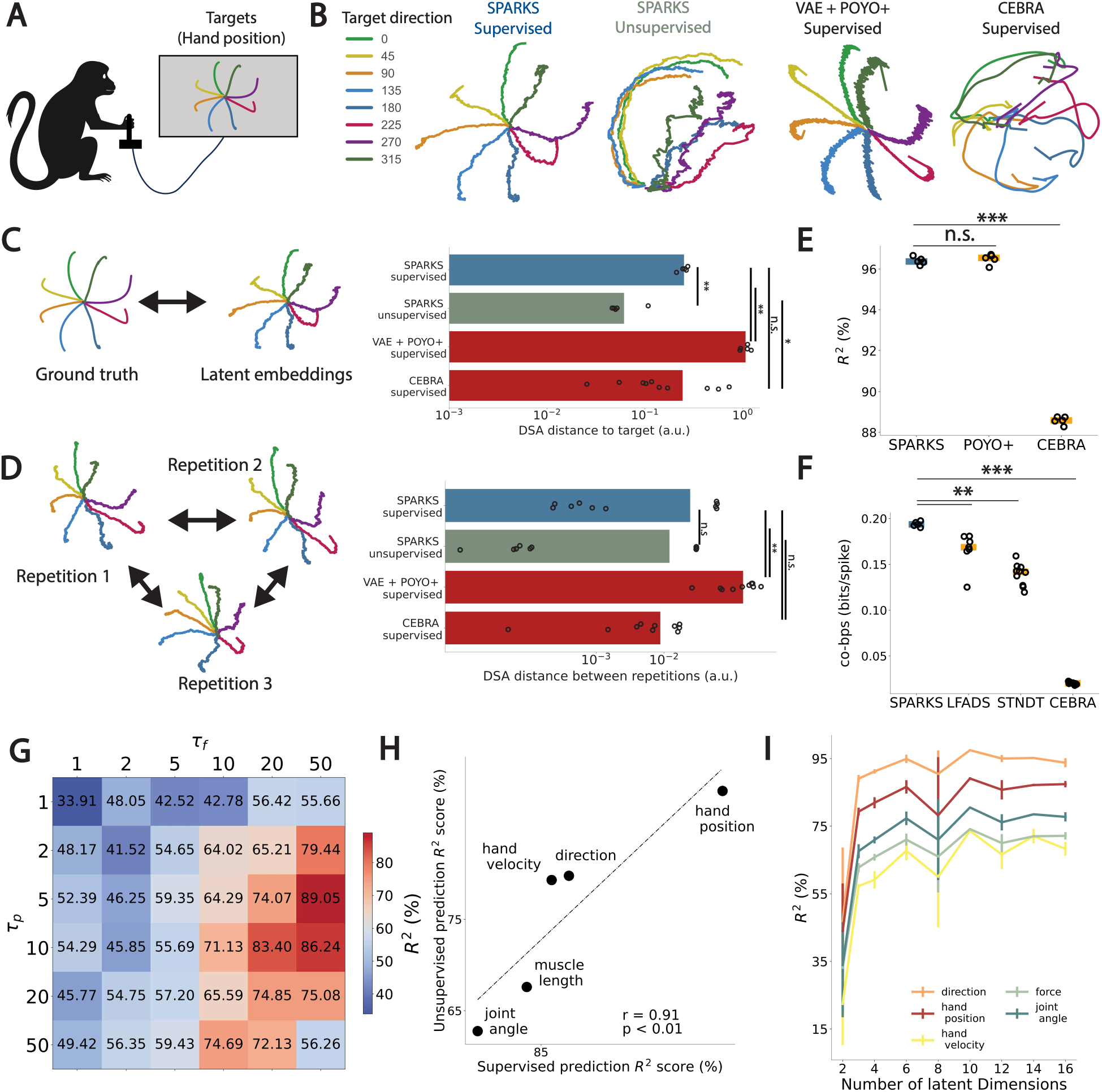
Dynamically faithful latent embeddings and state-of-the-art prediction from somatosensory cortex data. (**A**) Illustration of the reaching task with average hand positions across all target directions. (**B**) Average latent embeddings obtained through SPARKS-fixed for supervised and unsupervised learning, a VAE-Transformer hybrid with POYO+’s attention mechanism^21^, and CEBRA^42^. (**C**) Left: Illustration of the comparison between latent embeddings and reference signal. Right: Comparison of the DSA distances between latent embeddings and the task parameter for SPARKS-fixed and competing models. Each value represents the mean over 5 individual runs. (**D**) Left: Illustration of the comparison between latent embeddings across repetitions for SPARKS-fixed and competing models. Right: Comparison of the DSA distances between the latent embeddings across 5 repetitions after training with random weights initialisation for each model type. (**E**) Comparison of co-smoothing scores for prediction of left-out neurons for SPARKS-fixed and competing models. Orange lines represent the median over 5 runs, and each dot an individual run. (**F**) Comparison of co-smoothing and R^2^ scores for prediction of hand position for SPARKS-fixed and competing models. Coloured lines represent the median over 5 runs, and each dot an individual run. (**G**) Effect of the past and future windows τ_p_ and τ_f_ on prediction of the animal’s hand position from unsupervised latent embeddings for SPARKS-fixed. (**H**) Unsupervised vs. supervised prediction scores for the same target variables with SPARKS-fixed. (**I**) Effect of the number of latent dimensions on the prediction of the animal’s hand position from unsupervised latent embeddings with SPARKS-fixed. *: p < 0.05, **: p < 0.01, ***: p < 0.001, n.s.: not significant, two-sided Mann-Whitney U test. r: Pearson correlation coefficient. p: p-value.

To further discriminate between the latent embeddings produced by different models, we adopted the perspective that the brain implements a non-linear dynamical system, which our model approximates within the low-dimensional latent space learned. In particular, we relied on dynamical systems analysis^40^ (DSA), a method based on Koopman operator theory^41^ to compare the vector fields of dynamical systems. DSA relies on computing high-dimensional approximations of the actual Koopman operators to measure differences between dynamical systems via statistical shape analysis. We computed DSA distances between latent embeddings and our target signal (the trajectories of the hand), and found that SPARKS creates embeddings that are significantly more similar to this reference than all benchmarks we implemented (Figure 3C, Figure S3D). Importantly, although replacing Hebbian attention with various alternatives produced visually relevant results with low Procrustes distances, it did not recover the dynamics of the reference signal. Conversely, CEBRA embeddings were geometrically distant from the hand trajectories but comparatively close to them under DSA, showing that the two measures capture distinct properties of the latent space, and that only SPARKS performs well according to both. We also used DSA to compute the consistency of latent embeddings across different repetitions of the training procedure with random initialisations. The encodings obtained with SPARKS were competitive with benchmark models, both in the supervised and unsupervised modes (Figure 3D, Figure S3E).

Another key aspect of SPARKS is its high predictive power, which allowed us to obtain highly accurate reconstruction of task variables (Figure 3E). When training the model via supervised learning, predictive power was quantified as the *R*^2^ score obtained using examples held out for testing. This metric is directly obtained from the decoder’s outputs, and we obtain *R*^2^ = 96.4% for the decoding of hand position, a metric that remains reliable across a large range of hyperparameters (Figure S3F-G). SPARKS achieves predictive performance matching that of a state-of-the-art decoding model^20^ (*R*^2^ = 96.5%, *p* = 0.43, two-sided Mann-Whitney U-test), and surpassed dimensionality reduction models^17^ (*R*^2^ = 88.6%, *p* = 0.007), as well as all other benchmark models we implemented (Figure S3H). These results generalised to co-smoothing, where SPARKS outperforms models for neural data prediction^22^ and dimensionality reduction^17^ (Figure 3F). Our results reveal that predictive accuracy is not necessarily a sign of dynamical fidelity: while models optimised for decoding can achieve high predictive scores, we find that they fail to capture the underlying dynamics of the neural manifold (Figure 3C, E). In contrast, SPARKS’ unique ability to capture the co-variation of neurons at every time-instant in the recording allows it to follow the dynamics of the signal along the manifold^43^.

Tuning the past and future windows *τ*_*p*_ and *τ*_*f*_ had the largest effect when performing unsupervised learning, as measured by the decoding accuracy of the hand position from the learned latent embeddings using k-nearest neighbour regression (Figure 3G, supervised and co-smoothing reported in Figure S3F-G). We were able to decode other behavioural variables from the embeddings produced via unsupervised learning (Methods), and selected the five variables — hand position and velocity, target direction, moment of force applied to the controller, and muscle length — that exhibited the highest unsupervised accuracy. Training our model to decode each of these variables separately, we found that the scores obtained from supervised learning exhibit a remarkably high positive correlation with those obtained from unsupervised learning (Pearson correlation coefficient *r* = 0.91, *p* < 0.01, Figure 3H). The unsupervised model therefore encodes into the latent space information about the behavioural parameters found in the neural signal, weighted in accordance with their importance in the spike trains. Although obtaining three-dimensional latent embeddings is convenient for visual inspection, note that we do not claim here that this neuronal population’s responses are intrinsically embedded within a three-dimensional space. The population’s intrinsic dimensionality can be estimated by sweeping the model’s decoding performance of relevant parameters across latent dimensions and examining when this metric plateaus, which we found to be at 6 latent dimensions (Figure 3I, *p* < 0.05). Based on the estimated manifold, latent trajectories can be used to investigate biological questions by exploring how different latent dimensions relate to sensory, motor and state dependent signals.

Taken together, these results demonstrate how SPARKS produces meaningful and robust low-dimensional representations of neural data, providing state-of-the-art performance for supervised and unsupervised learning as well as generation of latent embeddings with only tens of neurons.

### Capturing Responses to Visual Stimuli Across Sessions and Animals

Next, we evaluated SPARKS’ capability to capture responses to higher-dimensional stimuli such as natural movies. To this end, we analysed Neuropixels recordings in the primary visual cortex obtained from the Allen visual coding dataset^44^, using passive recordings from mice viewing a 30 second natural movie (Figure 4A), and constructed a *pseudo-mouse* by aggregating the recordings from individual units, chosen at random across all 58 sessions, each recorded in a different animal.

**Figure 4.**
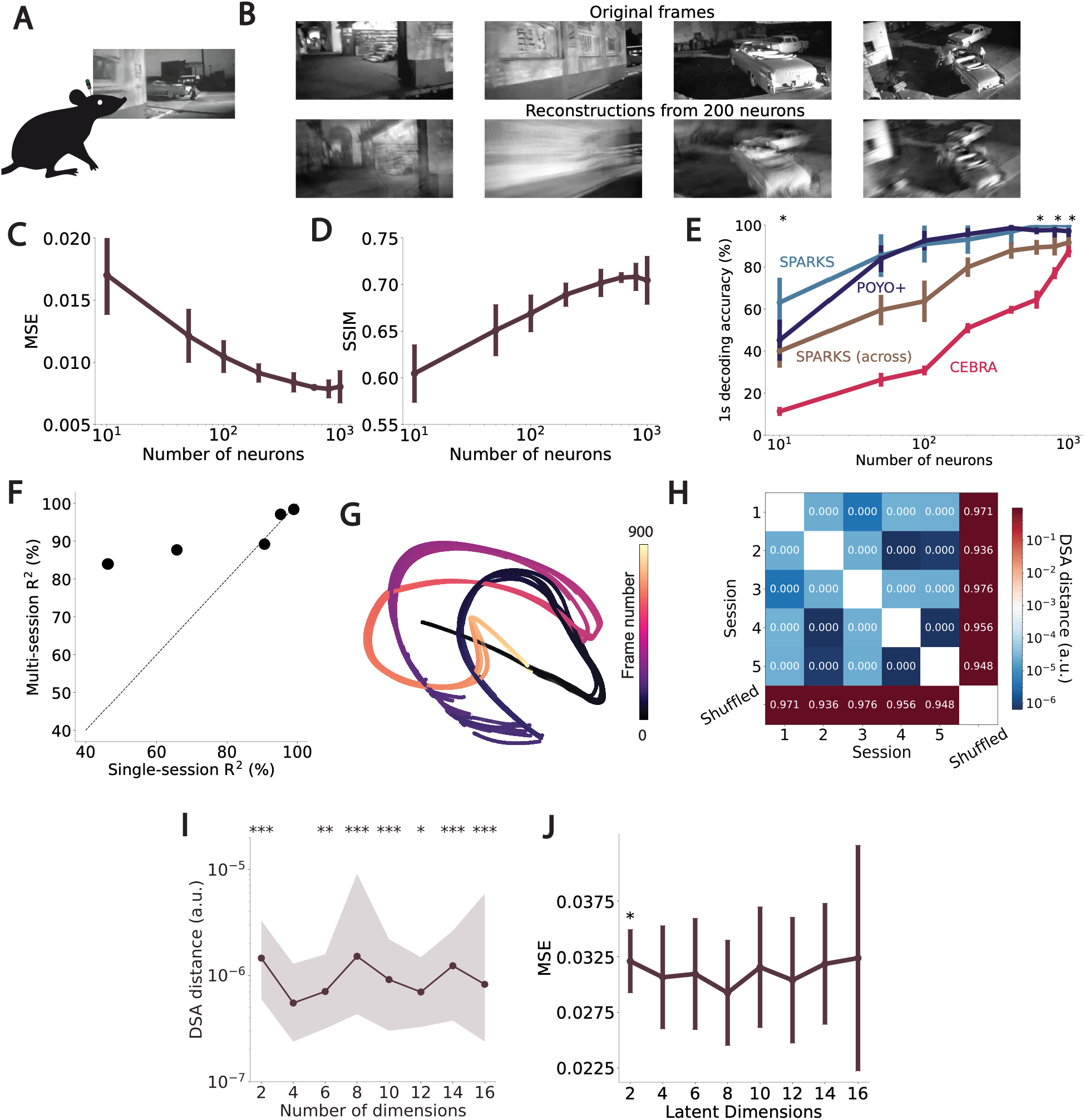
Frame reconstruction and prediction from primary visual cortex recordings. (**A**) Schematic of Neuropixels recordings in passive mice viewing a naturalistic film. (**B)** Example frames from the naturalistic movie and corresponding reconstructions using 200 neurons recorded from mouse primary visual cortex. (**C**) Reconstruction MSE as a function of number of neurons for SPARKS-fixed. Error bars represent standard deviations over ten repetitions with random sampling of individual units from all recordings. (**D**) Reconstruction SSIM as a function of number of neurons for SPARKS-fixed. Error bars represent standard deviations over ten repetitions with random sampling of individual units from all recordings. (**E**) 1s decoding accuracy as a function of number of neurons for SPARKS-learned and competing models. Error bars represent standard deviations over ten repetitions with random sampling of individual units from all recordings. Stars represent statistical difference between SPARKS’ and POYO+’s predictions, two-sided Mann-Whitney U test. All lines represent accuracy within one block of presentations, except SPARKS (across) that represents accuracy across the two blocks (Methods). (**F**) Scatter plot of the 1s prediction accuracy for the multi-session model as a function of the prediction accuracy of single-session SPARKS-fixed models. Each dot represents the performance for a single session, averaged over 5 repetitions. (**G**) Three-dimensional embeddings corresponding to different sessions obtained from the multi-session SPARKS-fixed model. (**H**) Pairwise DSA distances between latent embeddings obtained for all sessions from SPARKS-fixed for frames reconstruction. Shuffled: latent representations from session 0, randomly shuffled across the time dimension. (**I**) Average pairwise DSA distances between latent embeddings of five different sessions obtained with SPARKS-fixed as a function of the latent dimensionality. Shaded areas represent interquartile range across 5 repetitions and sessions. Stars represent statistical difference with the minimum obtained at 4 latent dimensions. (**J**) Average reconstruction MSE across sessions with SPARKS-fixed as a function of the latent dimensionality. Error bars represent standard deviation across repetitions and sessions. Stars represent statistical difference with the minimum obtained at 8 latent dimensions. *: p < 0.05, **: p < 0.01, *** : p < 0.001.

We started by reconstructing movie frames from neural data (Figure 4B). Reconstructions captured high-level scene context, as well as small, fast-moving details using only a few hundred neurons (Suppl. Movie 1). We measured the quality of reconstructions through mean-squared error (MSE) as well as structural similarity index measure (SSIM), a measure of perceived quality^45^. Both measures were seen to improve with the number of neurons aggregated, and saturated at around 600 neurons (Figure 4C-D). In this case, most alternative modules increased reconstruction MSE, whereas conventional attention reached an error comparable to SPARKS (Figure S4A). We also tested our model on the task of predicting the index of the frame currently shown to the animal (Figure 4E). SPARKS either matched or outperformed state-of-the-art decoding and dimensionality reduction models^21,42^, the latter using five times fewer neurons (CEBRA, 87.4% accuracy, *N* = 1,000; SPARKS, 93.03% accuracy, *N* = 200, *p* < 0.05). On this task, SPARKS-learned performed best, especially when the neuron count was high, suggesting that the benefits of this configuration can only be obtained if enough data is available to fit the extra learnable parameters (Figure S4B).

Movies were shown to the animals in two blocks of 10 presentations, separated by approximately 90 minutes. So far, we only discussed predictions formed using a single block of presentations of the same natural movie (Methods). Analysing neural activity recorded over periods ranging from tens of minutes to hours or more is known to be a difficult task due to the high level of variability across trials, which can be attributed to changes in neural state or behaviour, a phenomenon known as *neural drift*^46–49^. To assess whether SPARKS can isolate stimulus-relevant neural responses while ignoring sources of variability, we tackled the more challenging task of prediction *across* both blocks, and tested our model on the second set of 10 presentations after training on all examples from the first block. Remarkably, SPARKS maintains very high accuracy in this task (Figure 4E), even surpassing the performance of the reference dimensionality reduction model performing within-block decoding (*p* < 0.05 across all neuron counts). We also found the latent embeddings obtained from both blocks lie close within the latent space (Figure S4C), despite the fact that the model was trained on one block only and the large variability in the neural data across blocks, as we discuss in the next section.

Generalising neural signal analysis across longer time-scales (i.e., days) and animals is an even more challenging task, albeit a central requirement for any robust and comprehensive analysis. Despite recent methodological advances^5,50^, achieving stable chronic electrophysiological recordings in mice continues to be highly challenging due to technical limitations, and many studies still rely on acute recordings. Concatenating data from several recordings remains standard practice, particularly when only a limited number of neurons per session can be recorded^44,51^. However, this approach introduces additional complexity and can obscure meaningful variability in behaviour and internal state across trials and sessions, even when experimental conditions and stimulation protocols are tightly controlled^52,53^. Pooling data in this fashion is also generally impractical, especially in behavioural tasks where trial lengths vary due to differences in response times across animals. SPARKS can be used to overcome these difficulties by learning session-specific attention layers (Figure 1E) and keeping the rest of the architecture shared across sessions. Analysing new sessions from a previously trained model is also straightforward via the addition of a new Hebbian attention layer and finetuning. We find that our model reaches its final accuracy on the new session within ~50 finetuning epochs, with no loss of accuracy on the sessions it was pre-trained on (Figure S4D).

Our analyses can therefore be performed across sessions in a shared latent space. To illustrate how this approach is beneficial, we first separately trained models for frame prediction using data from 5 separate sessions, and found a high variance in the predictive accuracy across sessions (Figure 4F). This effect is not explained by the number of neurons in each recording (Figure S4E, *r* = 0.28, *p* > 0.1), suggesting that not all recordings from the same brain region are equally informative. However, when training a single model with all five sessions, we found a large improvement in prediction accuracy in sessions with low performance, that all latent representations lay close in the latent space, and that trajectories produced by the model remained highly consistent, as measured by DSA (Figure 4F-H, Figure S4F). This result points towards the existence of a shared stimulus-driven manifold, that can be recovered in sessions where prediction accuracy was low due to neural responses being dominated by e.g. behavioural responses.

SPARKS is therefore able to capture a shared manifold of neural activity across several recordings. Characterising such structures is an open question in neuroscience, as conflicting results suggest the intrinsic dimensionality of visual responses may lie anywhere from a handful of dimensions to the full neural state-space^54,55^. To estimate the dimensionality of this manifold, we concurrently monitored the reconstruction MSE and the pairwise DSA between the latent embeddings of all sessions for a range of latent dimensions (Figure 4I-J). We reasoned that as the number of latent dimensions increase, the capability of the model to reconstruct frames would also increase, and eventually saturate when the dimensionality of the recorded population’s visually-driven activity is reached. This should correspond to different values for each recording, as all are not equally informative about the movie (Figure 4G). Simultaneously, between-session DSA should decrease as the number of dimensions increase and the latent embeddings of each session capture more movie-driven signal, and therefore become more similar to each other. Pairwise DSA should reach a minimum at the dimension of the shared manifold, and subsequently increase, as additional dimensions will be used to fit more stimulus-driven representations in the informative sessions, but not in others. Interestingly, the MSE plateaued for all the sessions at the same point that the DSA started increasing, at 4 latent dimensions (Figure 4I-J). Because these two independent criteria identified the same dimensionality, we interpret the shared manifold as the manifold of visually-driven responses common to all sessions. As a control, we repeated the analysis in the unsupervised setting, where initial latent dimensions will carry the largest amount of variance, rather than stimulus relevance (Figure S4G-H). In this situation, the average pairwise DSA showed no trend up to 16 dimensions and the reconstruction MSE kept improving up to 10 dimensions, consistent with the leading dimensions in cortex carrying behaviour-related variance that is largely uninformative about the stimulus^6^. The MSE plateau coinciding with the DSA minimum is therefore specific to the shared stimulus-driven manifold, and not a generic consequence of adding dimensions. This preliminary result is based on small neurons counts and few sessions, warranting a detailed analysis that is out of the scope of this paper. It demonstrates, however, that when used appropriately, SPARKS offers a path to studying neural manifolds across sessions and animals.

### Isolating the Neural Code of Visual Information

In the previous section, we found that in a prediction task, SPARKS can ignore neural variability over large time-scales to capture the manifold of stimulus-driven responses. Isolating the reliable sensory responses encoded in a neural population is an open challenge in neuroscience research, and a key step towards understanding how populations of neurons encode information. To further investigate the properties of stimulus-driven neuronal responses in the Allen visual coding dataset, we employed SPARKS as a *denoising autoencoder*^56^ to segregate the fraction of the neural responses driven by the movie shown across the two blocks. Specifically, we trained our model by using as reference signal the neural sequence from another randomly selected movie presentation (Methods), forcing it to learn the common visual manifold across all trials (Figure 5B). After training, we found that SPARKS had learned common spiking patterns across all presentations of the movie, as demonstrated by the very high level of correlation between the predicted spiking probabilities for all neurons within, but also across blocks (Figure 5C). Because this training scheme explicitly favours trial-invariant features, we compared SPARKS with trial-averaged responses, the standard estimate of stimulus-evoked activity. Importantly, the correlations were not strictly 1, as one would expect from using averaged responses, showing that SPARKS can still capture on-manifold, per-presentation variability in the neural responses. Whereas a trial average assumes a fixed stimulus response across presentations, SPARKS re-estimates it on every trial, and we found that SPARKS predictions capture single-trial responses far better than trial averages (Figure S5A, median correlations: block 1 – SPARKS: 0.97, trial-averaged: 0.39, p < 1e − 10; block 2 – SPARKS: 0.97, trial-averaged: 0.39, p < 1e − 10). Therefore, while trial averaging fails under representational drift across the 90-minute block gap, SPARKS is able to re-estimate the stimulus response for every trial.

**Figure 5.**
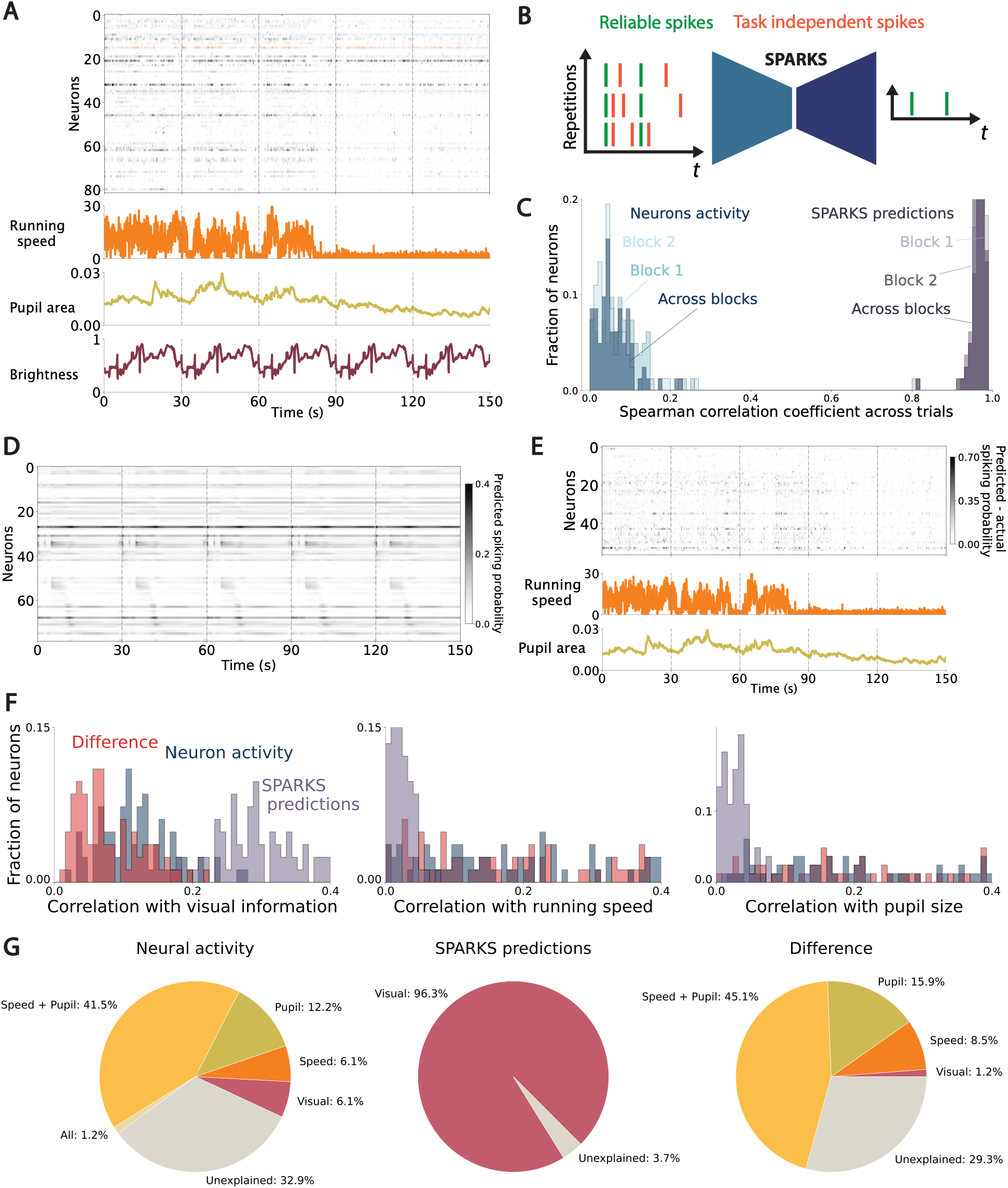
Isolating stimulus-driven and behaviour-related neural variance using SPARKS. (**A**) Representative activity of a population of neurons from the primary visual cortex of a passive mouse during presentation of a natural film; running speed, pupil area and average brightness of the 50 pixels most correlated to the neuron whose responses are highlighted in red on the top panel are shown below. Vertical lines correspond to the start of a presentation of the natural movie. Bars indicate firing events. (**B**) Illustration of SPARKS as a denoising autoencoder: across repetitions, neural responses present similar patterns that can be learned by the model. (**C**) Spearman correlation of the neurons’ responses across trial within each block, and across blocks. (**D**) Raster plot of predicted spiking probabilities with SPARKS-learned, for neurons in which predictions are significantly correlated to pixels brightness. Vertical lines correspond to the start of a presentation of the natural movie. (**E**) Top: Raster plot of the difference between neuronal activity and predictions with SPARKS-learned, for neurons in which the difference is significantly correlated to running, pupil area or both. Middle: running speed. Bottom pupil area for the same animal. Vertical lines correspond to the start of a presentation of the natural movie. (**F**) Distribution of correlation coefficients for raw neuronal activity (blue), SPARKS-learned predictions (purple), and for the difference between the two (pink) with pixels brightness (left), running speed (centre), and pupil size (right). (**G**) Pie charts representing the fractions of neuron correlated to measured parameters, based on their activity (left), SPARKS-learned predictions (centre), and the difference between the two (right).

SPARKS’ predictions strongly correlated to the brightness of pixels on the screen, which we use here as a proxy for visual information, creating repetitive spiking patterns across repetitions of the movie (Figure 5D). By looking at the difference between actual and predicted spikes, we were able to examine the variable part of the neural responses. For many neurons, the residual signal was correlated to the running speed of the animal and pupil size (Figure 5E and Figure 5F centre and right columns), but seldom to pixel intensities (Figure 5F, left column). Although a small (6.1%) fraction of neural responses were correlated to visual information only, we found that the largest group (41.5%) of neurons were correlated to a combination of speed and pupil size, with many cells (32.9%) presenting no correlation to a parameter recorded in this dataset (Figure 5G).

Remarkably, after denoising, the activity of nearly all (96.3%) neurons was correlated with visual content, revealing visual responses that were initially masked by behavioural variance. Trial-averaged responses recovered a smaller fraction (62.2%) of visually responsive neurons (Figure S5B-C), and the remaining neurons’ responses did not correlate to any parameter recorded. These results demonstrate SPARKS’ capability to separate the responses of neurons to various relevant parameters, offering a solution to the key problem of decomposing their variance into sensory responses and behavioural or state-related modulations.

### Uncovering Functional Interactions with Hebbian Attention

One of the benefits of biologically-inspired models such as SPARKS is their increased interpretability, as compared to black-box deep learning models. In our case, the Hebbian attention layer can be used to reveal biological insights in large neuronal networks. By capturing neuron-to-neuron interactions, we postulated that the coefficients calculated by the Hebbian attention layer could be reflective of functional interactions. To test this, we generated 6,720 hierarchical leaky integrate-and-fire networks with known connectivity (Methods). Networks comprised 3 to 6 modules. Within each module, we fixed the connection density and E/I balance, with each neuron following Dale’s law. Inter-areal connectivity was hierarchical and asymmetric: feedforward projections were both more numerous and stronger than feedback projections. The network was driven by independent Poisson input to every neuron in the lowest module in the hierarchy only (“thalamic drive”); to all modules (“uniform drive”); or to all modules, but with a stronger drive to the lowest module (“mixed drive”). We investigated whether we could recover the direction of the inter-areal hierarchy known by construction with SPARKS for a range of projections strengths and input firing rates (Methods). We found that by averaging the STDP coefficients calculated by Hebbian attention for all pairs of neurons from area *i* to area *j*, the resulting value could be used to estimate the flow from *i* to *j*, and use the total net outgoing flow per area to estimate the functional hierarchy (Methods). The hierarchy scores we recovered over all the configurations we tested strongly correlated with the actual, known hierarchies (Figure 6B, Figure S6D). We compared our predictions with the standard evaluation of functional connectivity based on cross-correlograms^57^ (CCGs, Figure S6E), and found that for all of the configurations we generated, the hierarchies predicted by SPARKS were equally or more correlated with the actual hierarchies than those predicted by CCGs (Table S2). With SPARKS, we were able to predict the correct position of modules in the hierarchy in 72% of cases (from 62% for six modules to 85% for three modules, Table S3), and our predictions were most accurate for modules at the top and at the bottom of the hierarchy (Table S4). For both metrics, SPARKS outperformed CCG-based hierarchies in most cases, with the exception of the least biologically realistic regime of a uniform drive (Tables S3–4).

**Figure 6.**
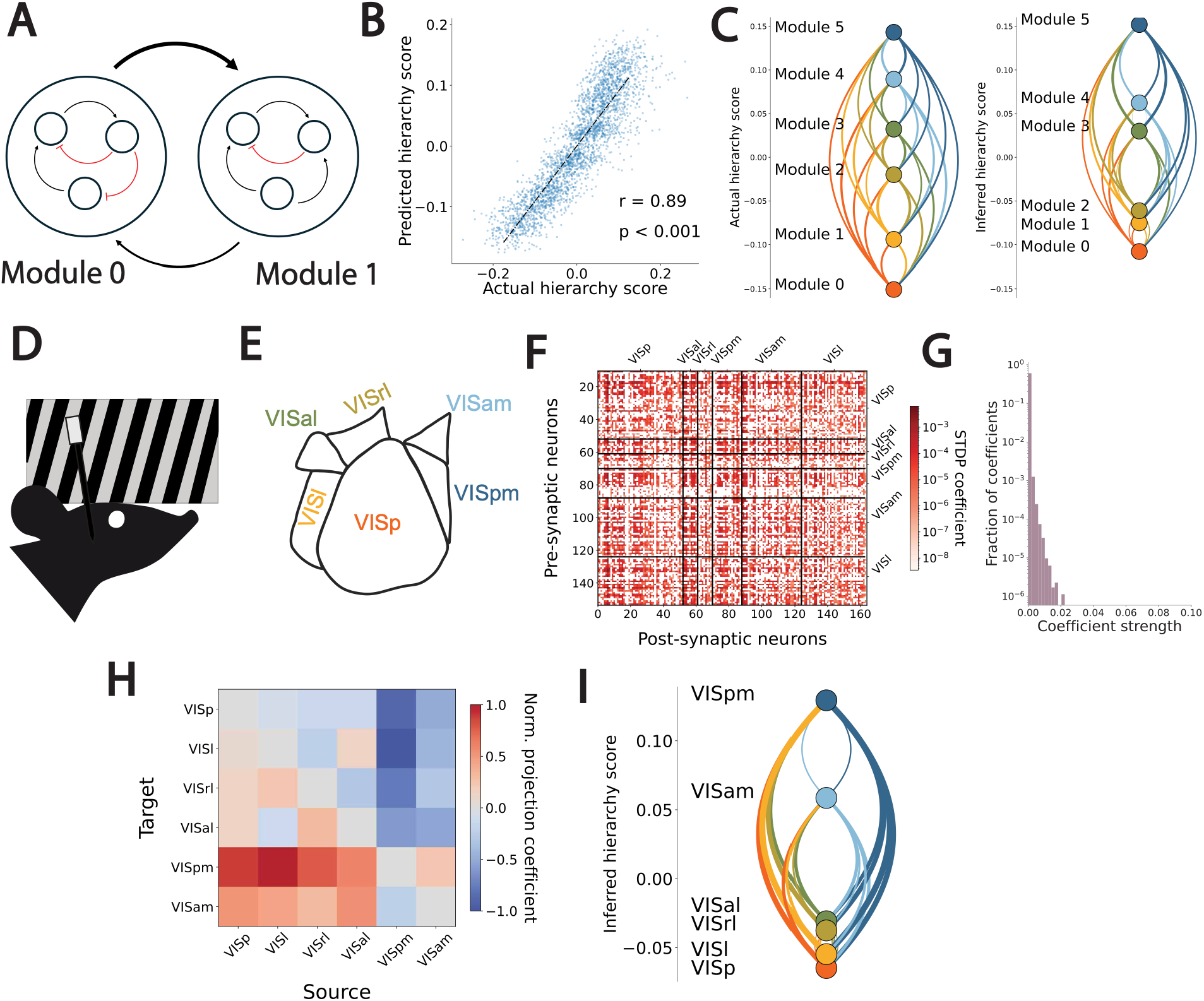
SPARKS recovers the hierarchy of the visual cortex. (**A**) Illustration of a hierarchical synthetic network of leaky-integrate-and-fire neurons with two modules. Black arrows: excitatory connections, line thickness illustrates relative connection strengths. Red arrows: inhibitory connections. (**B**) Correlation between the inferred and actual hierarchy scores for networks with six modules. (**C**) Example hierarchy based on a simulated network’s actual connectivity (left), and inferred hierarchy recovered with SPARKS-fixed (right). (**D**) Schematic of Neuropixels recordings in passive mice viewing a static grating. (**E**) Anatomical organisation of mouse visual cortical areas (**F**) Example STDP coefficients for one session from SPARKS-learned after subtraction of the shuffle baseline, on a logarithmic colour scale. Near-zero entries correspond to pairs whose coefficients do not exceed those expected from firing rates alone; area-level structure emerges only after pooling across pairs and sessions (**G**) Distribution of all STDP coefficients computed by SPARKS-learned as part of the Hebbian attention layer for 31 sessions (**H**) Projection scores between all visual cortical areas. (**I**) Functional hierarchy obtained with SPARKS-learned, calculated by computing the difference between their incoming and outgoing projections.

In the synthetic case, the model still had access to a perfect recording of all the neurons in the networks we generated. We investigated whether these results generalise to multi-region interactions using Neuropixels recordings from the Allen visual coding dataset^57^ acquired in the mouse primary visual cortex (VISp) and five higher-order visual cortical areas (HVAs): latero-medial (LM), antero-lateral (AL), rostro-lateral (RL), postero-medial (PM) and anteromedial (AM, Figure 6E). We trained SPARKS using neurons recorded in each HVA from awake, head-fixed mice passively viewing drifting gratings, static gratings and flashes. To limit the formation of possibly spurious correlations when collating neural responses from different sessions, we trained SPARKS using separate Hebbian attention heads for each of all 31 recordings, totalling 7080 neurons (Figure S6F). We found that the distribution of the STDP coefficients calculated by the model (Figure 6F) followed a similar distribution to those observed in experimental brain connectivity data, with most neuron pairs forming no connections, and presented a heavy tailed distribution (Figure 6G). Similarly to previous works^57,58^, we found that VISp is the area with the most outgoing projections, while areas VISam and VISpm receive the most inputs from other regions (Methods, Figure 6H).

We then computed hierarchy scores for each area, defined as the difference between incoming projections to the area and outgoing projections from that area. The functional hierarchy we obtained (Figure 6I) closely matched the anatomical and functional organisation of mouse cortical visual areas previously described^57,59^, strongly correlating with published hierarchy orders (Spearman *r* = 0.94, *p* = 0.005, significance evaluated by session-level bootstrapping, see Methods). This result can be understood by considering that the STDP coefficients inherently capture directed, temporal relationships between neurons. Therefore, they can be interpreted as a measure of an information flow, which serves as a proxy for the actual functional information flow in the biological circuit, demonstrating the interpretability of SPARKS’ Hebbian attention layer to reveal large-scale network interactions.

## Discussion

Motivated by the challenges arising from the collection of large-scale neuronal recordings and the surge of generative machine learning techniques, we developed SPARKS, a model combining the benefits of variational autoencoders and Transformers to obtain low-dimensional representations of neural data and very high predictive accuracy. Unlike traditional methods that rely on heavily processed firing rates, SPARKS operates at the single-spike level, capturing the temporal richness of neuronal responses while maintaining interpretability and scalability.

The Hebbian attention layer is the core innovation in SPARKS’ architecture, adapting the highly effective attention mechanism of Transformers to process spiking signals. It repurposes the mechanism of STDP into a spatio-temporal operator that is naturally suited to neuronal recordings. Hebbian attention enables our model to process neuronal information more effectively, offering state-of-the-art performance in various tasks and an increase in interpretability as compared to black-box deep learning models. Secondly, we introduce a novel learning objective based on predictive causally conditioned distributions and inspired by predictive coding in the brain. By forcing our model to predict the future evolution of the neural signal, the PCC informs the creation of highly reliable latent embeddings. This is a different approach to the currently popular paradigm of processing time-series data relying on contrastive learning^17^, where representations of “positive pairs” are pulled together, while those of “negative pairs” are pushed apart. The PCC objective can be understood as a generative analogue to these contrastive approaches, by constructing an explicit model of the data density relying on the assumption that neural and behavioural states evolve smoothly over short timescales. This objective can be applied to any latent variable model of neural data, in line with recent work highlighting the benefits for training LLMs to simultaneously predict several tokens^32^. A better understanding of this mechanism and of its links with predictive coding in biological brains will open new perspectives to improve the training of large-scale AI models. Our approach is inspired by the recent successes of physics- and biologically-inspired AI models^26^, and demonstrates the relevance of using synergies between AI systems and our biological understanding of intelligence.

As we have highlighted throughout, SPARKS can be used to investigate biological questions using three mechanisms. The first is its capability to discover low-dimensional representations of the high-dimensional neuronal data, or neural manifolds. The neural manifolds approach has been gaining traction since early works in the 2000s demonstrating that “populations are more than the sum of their neurons”, that is, the collective firing patterns of population of neurons exhibit properties that cannot be reduced to a simple linear superposition of their firing rates^14,15^. Recent work notably demonstrated that cortical population activity is resilient to the loss of single neurons, and adapted to maintain its manifolds, demonstrating the relevance of studying neural responses at this level^60^. SPARKS identifies neural manifolds by comparing, pair by pair, the timing of spikes emitted by all recorded neurons. Understanding the structure of neural manifolds can shed light on the algorithms the brain implements; a now classical example being the ring attractor topology of head direction in the drosophila^61^. Another approach involves studying the trajectories of neural responses in lower-dimensional spaces, for instance to evaluate how they differ across different sensory conditions by projecting them along different *coding directions*. The main limitation to this approach now lies in our restricted understanding of the mechanisms of generative AI. In this work, we chose to study the latent neural manifolds produced by SPARKS under the perspective of dynamical systems, acknowledging that they cannot reliably be understood under a traditional Euclidean perspective. This offers a theoretical grounding to the analysis of our latent spaces, through which we find that our model is highly reliable, despite the variability inherent to generative models. This angle, that we aim to pursue in future work, notably aligns with recent calls for an algorithmic understanding of generative AI^62^, and we propose that biologically-inspired models offer an avenue towards higher model interpretability. In SPARKS, we found that the STDP coefficients obtained within the Hebbian attention layer can be used to uncover the functional structure of large-scale neuronal populations, offering a second approach to analyse data. These coefficients are interpreted at the level of populations rather than individual synapses: with only a fraction of the circuit recorded, pairwise statistics are dominated by shared input and polysynaptic pathways, which bias connectivity inferred from activity^63^. This approach comes as an alternative to existing methods such as Granger causality^64^ (GC), where SPARKS provides an indirect, model-based inference of hierarchy, whereas GC is a direct statistical model of causality.

A final, complementary approach relies on supervised learning to predict relevant parameters such as behavioural outputs or sensory information. SPARKS’ state-of-the-art prediction capabilities can find relationships between neural activity and behaviour that previous models could not capture. In addition, our model’s structure opens new possibilities to solve challenging non-linear tasks, such as segregating relevant neural information from behavioural variance. Note that this approach differs from disentangled representation learning^25,65,66^ in that it does not aim to learn individual subspaces corresponding to distinct, interpretable factors of variation in the data. Rather, it focuses on isolating the stable, stimulus-driven parts of neural responses, with the benefit of not requiring explicit labels for behavioural variables during training. Importantly, since disentangled representation learning relies on VAEs, both approaches could be combined to yield greater benefits. Although predictive power was not our main goal, SPARKS matches or exceeds specialised decoders across datasets, suggesting that Hebbian attention is well suited to neural data and could serve as a building block for foundation models of the brain. These models, comprising many billions of parameters and trained on large quantities of data, are expected to provide novel insights on brain function thanks to their *emergent* properties – that is, the capacity to perform tasks they were not specifically trained on.

Our modelling can also be seen under the perspective of the Bayesian brain hypothesis, by considering SPARKS as a Bayesian encoding-decoding machine^67^. The neural signal measured and forming the inputs ***x*** ~ *p*(***x***|***z, r***) to our model depends on both the animal’s internal generative model of the world *p*(***z***|***r***), and its sensory inputs ***r***. In our model, the encoder performs the approximate posterior inference *p*_*w*_(***z***|***x, r***) given the neural response to the current sensory inputs and parametric weights ***w***. Due to the noise inherent to neural coding and the probabilistic nature of the encoding ***z***, the decoding network is a Bayesian decoder tasked to estimate *q*_*w*_(***r***|***z, x***). By training encoding and decoding networks together, we obtain latent embeddings that are highly consistent across animals and sessions, which aligns with the Bayesian brain hypothesis that the brain’s internal model is calibrated through evolution and exposure to natural stimuli, and evolves slowly during adulthood. This approach differs from existing unified encoding-decoding models^24^ that rely on purely discriminative masked autoencoders, and instead offers an unsupervised generative model to explore the data. SPARKS is also closely related to LFADS, another sequential variational autoencoder. Unlike LFADS, whose bidirectional encoders infer latents from the whole trial, the SPARKS encoder is causal and operates on spike timing through Hebbian attention; it is trained with the PCC rather than a trial-level ELBO, and handles multiple sessions through session-specific Hebbian layers.

Despite these advances, important challenges remain. In its learned configuration, which is better suited to larger-scale recordings as we show on the Allen visual coding dataset, the quadratic complexity of attention limits scalability, and our current implementation cannot yet handle the hundreds of thousands or more neurons recorded simultaneously. Future work will explore sparse attention mechanisms^68^ and the integration of anatomical connectivity to align model structure with the brain’s wiring constraints. Additionally, while SPARKS provides compelling insights into latent organisation, understanding how these representations map onto circuit mechanisms and behaviour will require targeted experimental validation. Addressing these challenges will not only enhance SPARKS but also inform the development of next-generation brain-inspired AI systems.

In conclusion, SPARKS marks a step toward a unified computational framework for neural data analysis, combining dimensionality reduction, predictive power, and biological interpretability. By integrating Hebbian attention with a predictive coding-inspired objective in a hybrid VAE– Transformer architecture, it matches or surpasses specialised state-of-the-art methods in diverse tasks — from decoding behaviour to revealing large-scale functional organisation. As datasets grow in scale and complexity, versatile and scalable tools like SPARKS will be essential for translating neural activity into insights about how the brain represents and predicts the world.

## Acknowledgments

The authors wish to thank O. Simeone, A. Schaefer, S. Tootoonian, M. Kollo and S. Gonzalo-Cogno for their discussions and comments on the manuscript, and members of the neuronal circuits and behaviour laboratory for their comments and suggestions on the figures.

## Funding

Biotechnology and Biological Sciences Research Council (BBSRC) BB/X013472/1

Francis Crick Institute CC2118

Wellcome Trust 225412/Z/22/Z

## Author contributions

Conceptualization: NS

Methodology: NS, FI, SS

Investigation: NS, NG

Visualization: NS

Data Acquisition: AEW

Funding acquisition: FI, SS

Project administration: FI, SS

Supervision: FI, SS, NS

Writing – original draft: NS, FI, SS, NG

Writing – review & editing: NS, FI, SS

## Data and materials availability

Main code repository: https://www.github.com/Iacaruso-lab/sparks. All data in the main text or supplementary materials are open source. Sources: Medial entorhinal cortex calcium imaging: https://github.com/soledadgcogno/Ultraslow-oscillatory-sequences/raw/main/Selection%20of%0codes/calcium_activity_matrix_60584_session17.mat, Area2_bump dataset from the Neural latents benchmark: https://neurallatents.github.io/, Allen visual coding dataset: https://observatory.brain-map.org/visualcoding. All scripts to generate synthetic data and reproduce results shown in this work can be found at https://www.github.com/Iacaruso-lab/sparks-examples.

## Methods

### Notations

For any two jointly distributed random vectors ***a*** = (***a***_1_, …, ***a***_*t*_) and *b* = (***b***_1_, …, ***b***_*t*_) from time 1 to *T*, at every time-instant *t*, we introduce the *predictive causally conditioned (PCC)* distribution of ***a*** given ***b*** as

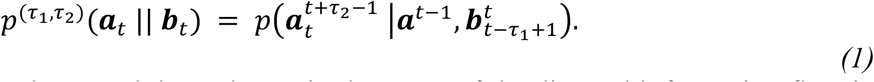

This distribution captures the causal dependence, in the sense of the directed information flow in time, of the *τ*_2_ future samples of RV 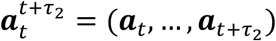 on the past *τ*_1_ samples of RV 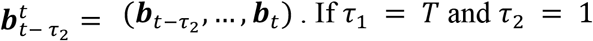, this recovers the causally conditioned distribution^29^.

### Problem definition

We study the autoencoder architecture in Figure 1A with the aim of training an encoding network to produce a representation ***z*** of the neural input ***x*** that is informative about a reference signal ***r***. The input ***x*** is a neural signal obtained using e.g. electrophysiology or imaging techniques. The reference signal ***r*** represents the corresponding target, such as the label corresponding to the behaviour of interest; a set of behavioural variables; or the input itself for unsupervised learning.

To simultaneously optimise encoder and decoder, we consider a data set comprising pairs (***x, r***) of neural inputs ***x*** and reference signals ***r***, jointly generated from an unknown population distribution *p*(***x, r***). The inputs are in the form of a discrete collection of vectors over time ***x*** = (***x***_1_, …, ***x***_*t*_) with ***x***_*t*_ ∈ ℝ^*N*^, and the reference signals ***r*** = (***r***_1_, …, ***r***_*t*_), with 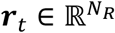, define general target signals, where we assume that all signals are uniformly sampled with a fixed sampling rate *dt*.

As seen in Figure 1A, the latent representation ***z*** = (***z***_1_, …, ***z***_*t*_) with 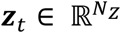 is obtained by processing the input signal ***x*** through the Hebbian encoder introduced in the main text and for which we provide mathematical details in the following. Given ***z***, the system is trained to predict signal ***r*** using a decoding network, implemented as an ANN. Following ideas from predictive coding, we maximise the causally conditioned distribution of *τ*_*f*_ future samples from the reference signal 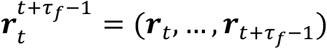 given past input spikes ***x***^*t*^ = (***x***_1_, …, ***x***_*t*_), averaged over time-steps *t* = 1, …, *T* as

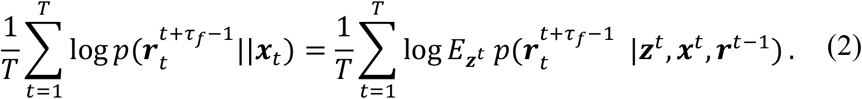

Note that the evaluation of objective (2) is intractable for problems of practical size^69^, and, as we detail in the following sections, we optimise over the model weights by considering a variational approximation developed in the following sections.

A single model may be trained jointly on multiple recordings from several sessions or animals, each with its own number of recorded neurons. In this case the problem above is instantiated once per recording.

### Encoding Network

Given the architecture proposed in Figure 1A, the encoder implements a causal stochastic mapping between input ***x*** and latent signal ***z*** that depends on a vector of parameters ***w***^*e*^ as

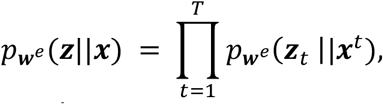

where the distribution 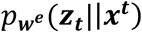 is obtained through the reparameterisation trick as

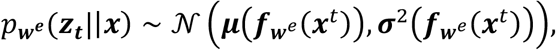

with 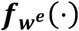 being the output of the encoding network. Samples are obtained as 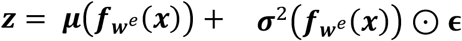 with ***ϵ*** ~ *N*(0, 1). The latent signal ***z*** is fed to a decoder, whose role is to predict the reference signal ***r***. Both blocks *µ*(⋅) and *σ*(⋅) are implemented as a single linear layer.

As we detail below, the encoding network 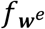 comprises, in order: a (session-specific) Hebbian attention block, which maps the *N*-dimensional input at each time-step onto a representation of dimension *M* × D; zero or more shared conventional attention blocks operating on that representation; and a final layer normalisation. When training the model over multiple sessions *i* = 1, …, *S*, we define session-specific parameters ***w***^*e*,(*i*)^, although we leave out the session index (*i*) in the following for notation clarity.

The optional conventional blocks implement causal dot-product self-attention^27^, with two modifications relative to the original description: positions are encoded by rotary position embeddings applied to the queries and keys^28^ in place of additive positional encodings, and each residual branch is scaled by a learnable per-channel gain initialised near zero^70^, so that a deep stack begins close to the identity mapping and the contribution of each block grows only as optimisation finds it useful. The default configuration uses no conventional blocks, in which case the Hebbian attention block alone constitutes 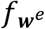.

The functions *µ*(⋅) and *σ*(⋅) are each implemented as a single linear layer reading off 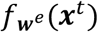. The log-variance is constrained to the interval [−4, 2] to add bounds on the magnitude of the injected reparameterisation noise. A layer normalisation is applied immediately before these two heads.

### Hebbian Attention

We propose to apply online STDP at every time-step *t* to compute a matrix ***A***_*t*_ ∈ ℝ^*N*×*N*^ in which each element *a*_*ij,t*_ defines a coefficient between sequences *i* and *j*. We consider an online STDP kernel, whereby coefficients for each synapse (*i* → *j*)_1≤*i,j*≤*N*_ are obtained by maintaining over time a pair of pre- and post-synaptic *eligibility traces* 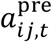 and 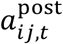 where *t* ∈ {1, …, *T*} and *N* is the number of recorded neurons. Without loss of generality, we assume *T* to be the (possibly infinite) temporal horizon of the task. When neurons on either side of the synapse fire, the corresponding traces are updated by an amount determined by (learnable) parameters 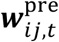 and 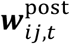, and exponentially decay in the absence of a spike by a value determined by (learnable) scalars 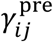 and 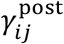.

The coefficient *a*_*ij,t*_ is potentiated (resp. depressed) by the value of the pre-(resp. post-) synaptic trace when the post-(resp. pre-) synaptic neuron fires.

At each time-step, the three quantities are updated as follows. The traces first decay,

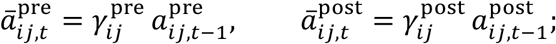

the coefficient is then updated using these decayed values,

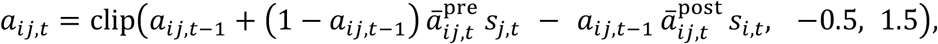

where *s*_*i,t*_ ∈ {0, 1} denotes the state of neuron *i* at time *t* and clip(⋅, −0.5, 1.5) is a hard bound enforcing its first argument to remain in the range [−0.5, 1.5]. Although the multiplicative factors (1 − *a*_*ij,t*−1_) and *a*_*ij,t*−1_ implement soft bounds that restrict coefficients to approximately the interval [0,1], we find that the clipping is necessary to prevent some coefficients from exploding when neurons exhibit high firing rates. STDP coefficients are initialised at zero at the start of a sequence.

Finally, the current time-step’s activity is incorporated into the traces

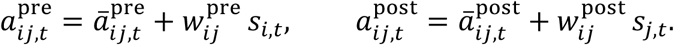

Note that the ordering of the above enforces the asymmetry of the rule: because coefficients *a*_*ij,t*_ are updated before the current spikes have been added to the eligibility traces, potentiation requires pre-synaptic activity that is strictly prior to a post-synaptic spike, and depression requires the converse; two neurons firing in the same bin therefore produce no potentiation. This asymmetry distinguishes the mechanism from generic co-activation measures.

### Parametrisation

The decay coefficients are defined through user-defined time constants 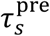 and 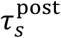 and the recording’s temporal resolution *dt* as

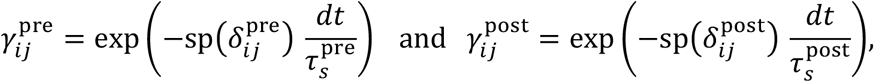

where unconstrained parameters 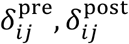 are initialised at sp^−1^(1) = log(*e* − 1), such that decay rates begin at exp(−*dt*/*τ*_*s*_) and the SoftPlus function sp(*x*) = ln(1 + exp *x*) operates in its approximately linear regime, where gradients are well conditioned.

Pre- and post-synaptic update weights are defined as

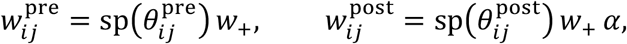

where *w*_+_ and *α* are fixed hyperparameters. The factor *α* > 1 biases the post-synaptic increment above the pre-synaptic one, breaking the symmetry between the potentiation and depression pathways at initialisation.

Two configurations of the layer differ in how these parameters are shared across pairs. In the learned configuration, 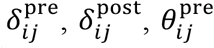 and 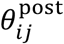 are learned independently for every ordered pair, amounting to 4*N*^2^ learnable parameters in the attention layer.

In the fixed configuration, these quantities are instead held fixed and shared across the population, so that *γ*^pre^ = *γ*^post^ = exp(−*dt*/*τ*_*s*_), *w*^pre^ = *w*_+_ and *w*^post^ = *w*_+_*α*, and the only learnable parameters of the attention layer are those of the learnable value projection. Therefore, the eligibility traces need only be maintained per neuron rather than per pair, reducing the recurrent state from *O*(*N*^2^) to *O*(*N*) until the coefficient matrix itself is formed at the combination step. Furthermore, training the model in this configuration does not require backpropagation-through-time.

Finally, the coefficient matrix is mapped onto a per-neuron embedding by an affine transformation applied along the post-synaptic index,

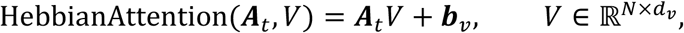

where ***b***_*v*_ is a learnable bias, so that the embedding of neuron *i* at time *t* is obtained from the *i*-th row of ***A***_*t*_, that is, from the coefficients relating neuron *i* to every other neuron. The matrix *V* is defined in analogy to the values of conventional dot-product attention^27^.

### Multi-headed attention

The mechanism above may be replicated to obtain several parameters sets (*T*^pre,(*k*)^, *T*^post,*k*)^, *W*^pre,(*k*)^, *W*^post,(*k*)^, *V*^(*k*^) for *k* = 1, …, *K*, each with its own trace time constant 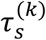 and therefore its own eligibility dynamics and value projection. The resulting multi-headed module allows the model to attend jointly to information from different representation subspaces^27^, and here specifically to correlation structure at several timescales simultaneously. Head outputs are concatenated and projected back onto *D* by a learnable matrix *W*_*o*_ ∈ ℝ^D×D^,

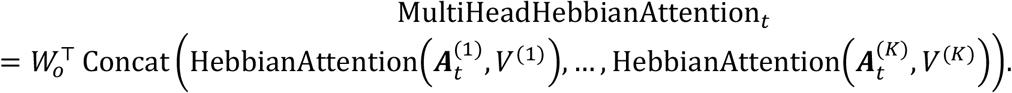

With *K* = 1 this reduces to the single-head mechanism above.

### Feedforward block

The embedding produced by the Hebbian attention layer is passed through a position-wise feedforward sublayer with a residual connection,

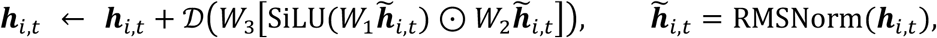

where the gated linear unit has inner width 8/3 *d*_*v*_ and no biases^71^, and *D*(⋅) denotes stochastic depth^72^, which independently drops the entire residual branch for each sequence in a mini-batch during training and rescales the surviving branches so that the expectation is unchanged. Normalisation is applied to the input of the sublayer rather than to its output, so that the residual path remains an exact identity mapping.

### Perceiver bottleneck

The *N* neuron-indexed embeddings are compressed onto *M* ≪ *N* latent slots by a cross-attention module in the manner of a spatial Perceiver architecture^73^.

A learnable embedding 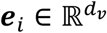, shared across time and specific to neuron *i*, is added before normalisation,

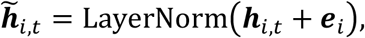

such that *M* learnable query vectors, which do not depend on the input and are shared across sessions, read out the population. The readout is implemented with a kernelised linear attention that carries a decaying recurrent state across time^74^. With the feature map *ϕ*(*u*) = elu(*u*) + 1, which guarantees positivity, applied to per-head projections 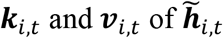,, the per-time-step contributions of the population are

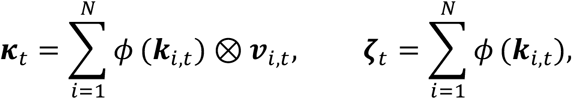

which are accumulated into a state that decays at a learnable, head-specific rate *λ*_*h*_ = sigmoid(l_*h*_) ∈ (0,1) with 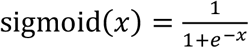,

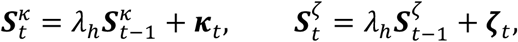

and read out by the *M* queries as

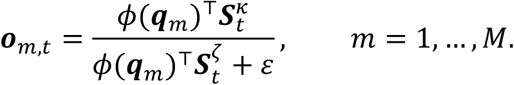

The rates *λ*_*h*_ are initialised at sigmoid(0) = 1/2, a neutral value from which optimisation may sharpen the state in either direction.

The *M* resulting slots subsequently attend to one another through multi-head self-attention, so that the compressed summaries are not independent, and pass through a further feedforward sublayer (Figure S1B). Both operations are applied with pre-normalised residual connections. The *M* slots are finally concatenated into a vector of dimension *M*D.

### Conventional layers

The concatenated outputs of the latent bottleneck are then flattened across the spatial dimensions, resulting in coefficients of size [*T, M*D], and passed to optional conventional blocks implementing causal dot-product self-attention^27^. Temporal positions are encoded by rotary position embeddings applied to the queries and keys^28^, and each residual branch is scaled by a learnable per-channel gain initialised near zero^70^, which we found important to stabilise training for deeper encoders.

### Calcium recordings

When considering calcium recording data, we propose to directly use the calcium traces of each neuron as eligibility traces, which removes the heavy processing that comes with spike deconvolution. Mathematically, given inputs ***x***, we have

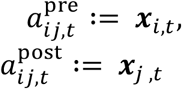

and

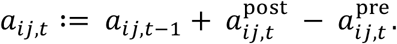

Even in this context, the Hebbian attention mechanism remains highly effective for two key reasons. Firstly, because the eligibility trace in our model is fundamentally a decaying memory of a past event, a raw calcium trace can be viewed as a biologically generated eligibility trace. The relative timing of the changes in these traces between neuron pairs provides a robust, albeit temporally smoothed, proxy for the precise spike-timing relationships that SPARKS is built to capture.

### Parallel prefix scan

Every recurrent quantity introduced above — the eligibility traces, the recurrent state of the linear-attention read-out, and the calcium traces, satisfy an affine recurrence of the form *x*_*t*_ = *a*_*t*_*x*_*t*−1_ + *b*_*t*_ in which *a*_*t*_ and *b*_*t*_ are known in advance of the recursion. The composition of affine maps being associative, such a recurrence admits an exact closed-form solution computable by a parallel prefix scan^75–77^ in *O*(log_2_*T*) sequential rounds of elementwise operations over the full sequence, in place of *T* strictly sequential steps.

The STDP coefficient update is the sole exception, as the hard bound breaks associativity. This recurrence is therefore evaluated sequentially, but with *a*_*t*_ and *b*_*t*_ precomputed for all *t* so that each step reduces to a single fused multiply–add–clip over the batch and pair dimensions, rather than a full trace-decay-and-update.

### Maximum likelihood optimisation

In order to address the maximisation of likelihood (2), we adopt a variational formulation that relies on the introduction of a decoding network that implements at every time-step *t* = 1, …, *T* the PCC distribution

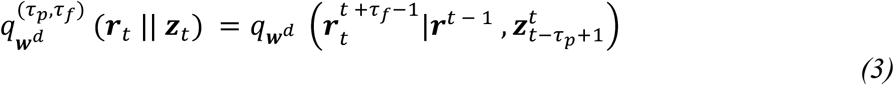

between latent representation ***z*** and reference signal ***r***^30,78^. The decoder *(3)* is causal, with memory given by integer *τ*_*p*_ > 1 and is parametrised by a vector ***w***^*d*^.

In practice, we implement the decoder as a model 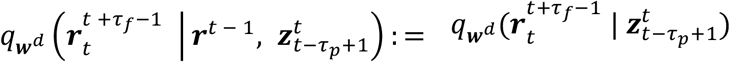, where 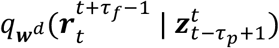 is implemented by the output layer of an ANN with inputs given by a window 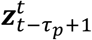 of samples from the latent representation ***z***. Alternatively, one can use an RNN or a probabilistic SNN ^79^ to directly model the kernel 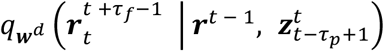.

With such a network, we use a standard variational inequality^69^ to lower bound the decoder log-likelihood at each step *t* = 1, …, *T*

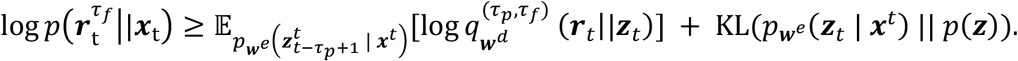

Averaging over time-steps, we obtain the *variational predictive likelihood*

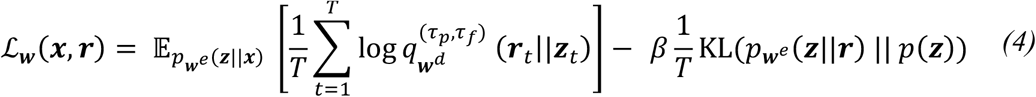

where ***w*** = (***w***^*e*^, ***w***^*d*^) and *β* is a hyperparameter. We maximise this objective via stochastic gradient descent (SGD) and the AdamW optimiser^80^ using Monte Carlo gradients. Note that with *τ*_*p*_ = 1 and *τ*_*f*_ = 1, the variational predictive likelihood reduces to the standard evidence lower bound (ELBO).

The gradient with respect to the encoder and decoder weights can be obtained via the stochastic gradient variational Bayes (SGVB) technique^81^ as

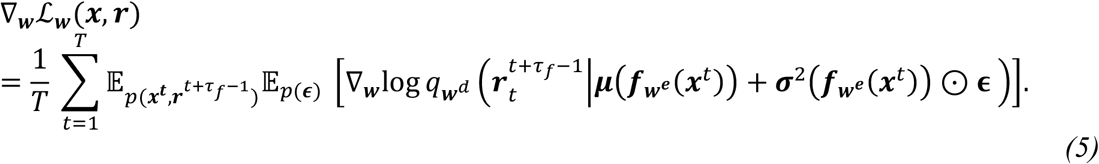

Unbiased estimates of the gradients can be evaluated by using mini-batches ℬ ⊆ *D* from the data set *D*, along with random samples 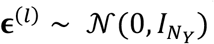 for 1 ≤ *l* ≤ *L* with *L* ≥ 1. The gradients **∇**_***w***_ w.r.t. the encoder and decoder weights are computed using backpropagation and automatic differentiation tools.

### Latent prior

Similarly to sequential latent-variable models for time series^82^, we implement a learnable autoregressive prior *p*(***z***_*t*_ ∣ ***z***_*t*−1_). A small, gated network predicts the prior from the sampled previous latent,

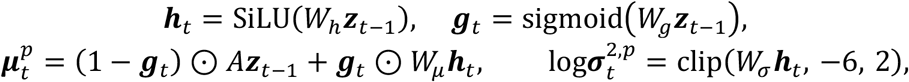

with *A* initialised to the identity, such that the transition begins as an approximately identity one-step-ahead predictor. The learnable per-dimension gate ***g***_*t*_ interpolates between this linear persistence prediction and a nonlinear correction. The corresponding divergence, for *t* ≥ 2, is evaluated per time-step and per dimension as

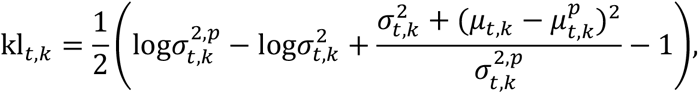

where the first time-step is initialised to a standard normal static regulariser, since no ***z***_0_ is available to condition upon. The parameters of the regulariser are appended to ***w*** and optimised jointly with the encoder and decoder.

### Truncated optimisation

The computation of gradients *(5)* is expensive as it requires to apply the backpropagation-through-time algorithm, which scales quadratically in terms of memory consumption with the number of time-steps. To alleviate memory issues during optimisation when the recording duration is large, a recording is divided into non-overlapping chunks of *C* time-steps, within which the forward dynamics of every recurrent component evolve without approximation, and the backward pass alone is truncated at chunk boundaries. The value of every recurrent state is carried from one chunk to the next, while its dependence on the preceding chunk’s computation graph is discarded, in the manner of truncated backpropagation through time^83^. Each chunk contributes an independent gradient estimate and parameter update, so memory consumption is independent of the number of chunks into which the recording is divided.

### Fixed vs. Learned configurations benchmarking

To characterise how the cost of training SPARKS scales with population size and recording length across both configurations, we trained the model via unsupervised learning on synthetic spike trains of *N* ∈ {10, 50, 100, 200, 400, 600, 800, 1,000} neurons and duration *T* ∈ {0.1, 0.2, 0.4, 0.6, 0.8, 1, 2, 4, 6, 8, 10} *s*, sampled at *dt* = 1 ms. Spikes were drawn independently for every neuron and time step from a Bernoulli distribution with probability 0.005 (5 Hz), so that cost depended only on *N* and *T* and not on the structure of the data.

The learned configuration was trained either by backpropagation through the whole sequence or by truncated backpropagation through time over consecutive chunks of 100, 500, 1,000 or 5,000 steps. The fixed configuration was trained without chunking. All other settings were shared across the fixed and learned configurations: embedding dimension 32, latent dimension 2, bottleneck dimension 8, *τ*_*p*_ = *τ*_*f*_ = 1, and linear decoder. Each of the combinations was run as a separate job on a single NVIDIA A100 SXM4 GPU (80 GB) for 20 epochs. For every epoch, we recorded the wall-clock training time, synchronising the GPU before and after the epoch, and the peak GPU memory allocated by PyTorch, reset at the start of each epoch. We report the mean and standard deviation of both quantities over all epochs except the first, which includes CUDA initialisation; runs that exceeded the GPU memory were recorded as failed.

Without chunking, peak memory grew linearly with recording length in both configurations as the model stores the whole sequence of *N* × *N* × *T* STDP coefficients in the Hebbian attention and subsequent layers perform causal attention across the time dimension. Chunking allows exact gradients calculation in the fixed configuration, but requires to re-allocate the tensor of STDP coefficients, which may be slower than performing one pass through the whole sequence, and should be tuned for each use case independently. Chunked training removed this dependence: peak memory varied by less than 5% across recording lengths from 0.2 to 10 s and was set by *N* and the chunk length alone. Runtime per epoch grew linearly with *T* under chunking. For populations of 400 neurons or more, shorter chunks reduced both memory and runtime.

### Alternative Self-Attention Mechanisms

To understand what components of the proposed Hebbian attention support our model’s performance, we substituted the corresponding block with each of the mechanisms described in this section, while leaving every other component of the architecture and of the objective unchanged.

The first two replace only the STDP computation in the Hebbian attention layer, retaining the coefficient-matrix formulation, and are routed through the same value projection, feedforward sublayer and latent bottleneck as our model; therefore, each isolated a single property of the coefficient computation at a time, namely the requirement for asymmetric Hebbian updates, and the requirement of single-spike resolution.

In the remaining three controls, we replace the whole attention block (including Hebbian attention layer, Feedforward and Perceiver blocks) to address the question of whether any explicit pairwise structure over neurons is required. We found that models in which we only replaced Hebbian attention with these modules did not converge during training.

Defining 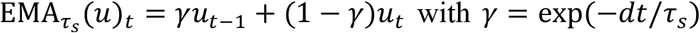 for the exponential moving average at timescale *τ*_*s*_, these three models map the smoothed population activity *ρ*_*i,t*_ directly onto an *Md*_*v*_-dimensional representation. They test whether explicit pairwise structure over neurons confers any advantage over generic sequence models operating on the same smoothed input under the same causal constraint.

#### Symmetric Hebbian attention

To determine whether the causal asymmetry of Hebbian attention is necessary, as opposed to Hebbian co-activation in general, we introduce a symmetric mechanism in which the coefficient is potentiated irrespective of firing order. The formulation of the original description retains per-pair traces and adds the two contributions symmetrically,

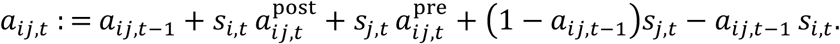

#### Instantaneous correlation

To determine whether pairwise, single-spike temporal dynamics are required, as opposed to a fixed short-memory statistic of neurons co-activation, we introduce a mechanism whose coefficients are a running, causal Pearson correlation between the smoothed rates of each pair of neurons.

The construction proceeds in three stages:

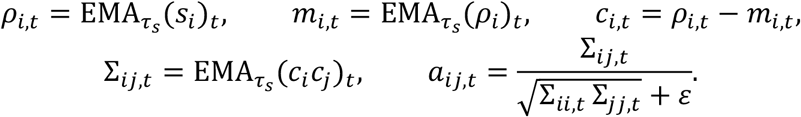

By applying a second EMA activation to the coefficients *ρ*_*i,t*_, we obtain smooth coefficients that are then centered and normalised. The unnormalised outer product *ρ*_*i,t*_*ρ*_*j,t*_, which is the immediate alternative, is non-negative everywhere and is dominated by shared population drive, so that any two neurons firing at high rates appear strongly related irrespective of whether their activity co-fluctuates; centring against the slower baseline *m*_i,t_ and normalising by the running variances yields a quantity confined to approximately [−1, 1] that reflects co-fluctuation.

#### MLP

We applied a position-wise multilayer perceptron with one hidden layer of width 4D and a Gaussian error linear unit nonlinearity, so that the only mixing across neurons is that effected by its dense weights and the only recurrence is that of the moving average itself.

#### GRU

The second applies a gated recurrent unit^84^ with hidden width D followed by a linear read-out, providing a recurrent baseline with a learnable hidden state but no attention.

#### Conventional causal self-attention

Finally, we applied the same causal self-attention block used in the conventional stack described above, preceded and followed by linear projections.

### Benchmark models

#### POYO+

We used the implementation provided by the authors at https://github.com/neuro-galaxy/torch_brain, and adapted the hyperparameters to match parameter counts between models (Table S1).

When replacing Hebbian attention with POYO+’s attention mechanism, we used the POYO+ encoder, to which we apply the reparameterisation trick to obtain latent embeddings.

#### STNDT

We used the implementation provided by the authors at https://github.com/trungle93/STNDT/tree/main and the parameters they provided where available, or the best parameters found after a search over 100 configurations with Ray Tune^85^.

#### LFADS

We used the implementation provided by the authors at https://github.com/arsedler9/lfads-torch/tree/main and parameters they provided where available, or the best parameters found after a search over 100 configurations with Ray Tune^85^.

### Model evaluation

#### Co-smoothing

For unsupervised learning, results were evaluated numerically through co-smoothing. This metric is based on the difference between the log-likelihood of a Poisson model with average firing rates ***λ*** corresponding to model predictions for left-out neurons outputs 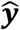, and a mean model with average value 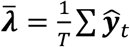 the mean firing rate of the neurons. Results are measured in bits/spike as

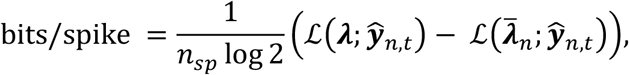

where *n*_*sp*_ is the total number of spikes.

#### DSA

We used the authors’ implementation at https://github.com/mitchellostrow/DSA. For each configuration, the number of delays and the rank of the Koopman approximation were selected by a parameter sweep, choosing the values that minimised the Akaike information criterion and mean absolute scaled error. When comparing configurations (e.g. across models in Fig. 3C or latent dimensionalities in Fig. 4I), the same parameters were used for all, set to the best fit across them.

### Configurations

Simulation code including all hyperparameters and model configurations for each panel in the main and supplementary figures can be found at https://github.com/Iacaruso-lab/sparks-examples.

### Mouse MEC dataset

Activity from MEC cells of head-fixed mice allowed to run on a rotating wheel in darkness was recorded using two-photon calcium imaging^35^. We considered a single example recording provided by the authors at https://github.com/soledadgcogno/Ultraslow-oscillatory-sequences/tree/main, which included 464 neurons recorded over 3600.51 seconds at a rate of 7.73 Hz. Following the authors’ methodology^35^, we downsampled the recording by a factor of 4 down to 1.93 Hz.

For the prediction task shown in Figure 2, we trained both SPARKS and another autoencoder with the same architecture but implementing the alternative symmetric Hebbian rule on the first 2600s of the recording, and leaving out the remaining 1000 seconds and 10% of randomly selected neurons for testing.

We show in Figure S2 the PSD of reconstructions, evaluated using Welch’s method with Hamming windows of 66 s (128 bins of 516 ms in each window) and 50% of overlap between consecutive windows.

From the 2-dimensional latent embeddings obtained, we recovered a phase signal by computing at every time-step *t* the angle between a segment joining the current embedding ***z***_*t*_ to the centre of the ring topology, and another joining the embedding ***z***_*t*_ to a fixed direction (here, the horizontal, Figure S2A). The phase signal thus obtained is shown in Figure S2B. For sorting with SPARKS, we sorted neurons by ordering them according to their correlation to this phase signal.

We evaluated the quality of each sorting by population decoding of the sorted signals (Figure S2C) and plotting their PSDs (Figure S2D). The population decoding 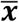 of spiking signal ***x*** was obtained as

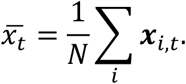

Similarly to the oscillatory score proposed in^35^, we evaluated the quality of each sorting by measuring the height of the peak above 0 in the PSD.

### Macaque reaching task dataset

The dataset corresponds to a single session extracted from^37^. Briefly, electrophysiological recordings were taken from Area 2 of the somatosensory cortex (S1) of a rhesus macaque during an eight-direction center-out reaching task. We considered a session obtained from the Neural Latents Benchmark^38^ to train our model. We used 1 ms bin times, and considered recordings from −100 ms to 500 ms after movement onset. Similarly to^38^, we considered a 40 ms lag between neural data and movement to align trials.

Results from CEBRA^17^ were shared by the authors at https://github.com/AdaptiveMotorControlLab/cebra-figures.

For prediction from the unsupervised latent embeddings, we first trained SPARKS via unsupervised learning using all examples, and then trained k-nearest neighbour regression models with 500 neighbours using a subset of examples. Training and test datasets for k-NN models were kept the same as for supervised learning with SPARKS.

A summary of parameters used for each model used in this section can be found in Table S1.

### Allen visual coding dataset

The Allen visual coding dataset comprises two-photon calcium imaging and Neuropixels data recorded from the primary visual cortex of mice during presentation of video scenes, and is available to download from the Allen Institute^44^. We only considered Neuropixels recordings. To construct a pseudo-mouse, we considered units sampled at random from all sessions in the Allen visual coding dataset, and stack the raw spike trains to perform movie reconstruction, frame prediction or unsupervised learning.

Each presentation of the excerpt from the movie *Touch of Evil*, shown at a rate of 30 Hz and lasting 30 s, was downsampled with a bin size of 5 ms. For frame reconstruction, movie frames *F* ∈ ℝ^900 × *h* × *w*^ were spatially downsampled to obtain film tensor 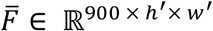 to reduce the computational load of the technique with h^′^ = h/2 and *w*^′^ = *w*/2.

The 1-second frame prediction accuracy was computed as

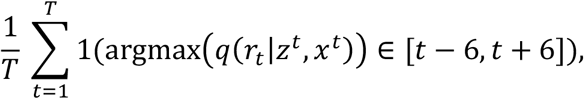

where 1(⋅) is the indicator function that outputs 1 if its argument is true, and 0 otherwise; and the argmax function outputs the largest index of its vector argument. The movie segment was presented 20 times in two blocks of 10 successive presentations, separated by roughly 90 minutes. Prediction within block was performed by training the model on the first 9 presentations in the first block, and testing on the last presentation. Prediction across block was performed by training on all examples from the first block, and testing on all examples in the second block. Reconstruction of frames in unsupervised learning was performed by training an auxiliary linear layer from latent embeddings.

#### Denoising

When using SPARKS for denoising, we used all 20 presentations of the movie 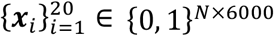 in both blocks, with *N* the number of neurons in the desired recording. At every epoch, we trained the model via self-supervised learning to minimise the loss

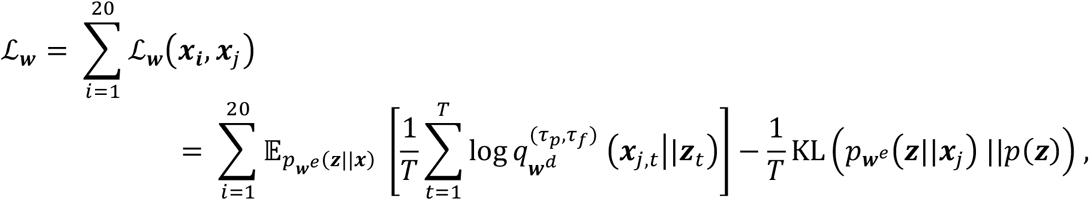

where the reference signal ***r*** for input sequence ***x***_***i***_ obtained during the *i*th presentation of the movie was selected as the input sequence ***x***_***j***_ obtained during the *j*th presentation, with *j* ≠ *i* selected at random.

Correlations with the neural signal were calculated after temporal smoothing obtained by convolving the neural recordings with a centered 50 ms boxcar filter (a moving average over 10 bins). Correlations were considered significant if the Spearman correlation coefficient was above the threshold *r* > 0.2.

### Synthetic hierarchical spiking networks

#### Network model

Ground-truth data was generated with recurrent networks of current-based leaky integrate-and-fire (LIF) neurons, implemented in PyTorch with snnTorch and simulated at dt = 1 ms. A network consisted of *B* modules of *N* = 100 neurons each; within every module the first 80% of neurons were excitatory and the remaining 20% inhibitory. Connectivity was split into a within-module matrix ***W***^loc^ and a between-module matrix ***W***^int^, both of size *BN* × *BN*, with the convention *W*_*ab*_ = weight of the synapse from presynaptic neuron *b* onto postsynaptic neuron *a*.

Within each module, connections were drawn independently for each ordered pair with probability *p*_*local*_ = 0.15, excluding self-connections, and nonzero weights were sampled from a Gamma distribution with shape 0.1 and rate 1. Dale’s law was imposed pre-synaptically: all outgoing weights of an inhibitory neuron were multiplied by −1, so inhibition was strictly local.

Between modules, the module index defined the position in the hierarchy. For an ordered pair (*m* → *m*′) with *m*′ > *m* (feedforward), synapses were drawn with probability *p*_*ff*_ and weights from *Gamma*(0.1, 1.0); for *m*′ < *m* (feedback), with probability *p*_*fb*_ < *p*_*ff*_ and weights from *Gamma*(0.1, 3.0), i.e. one third of the mean feedforward amplitude. Only excitatory neurons projected between modules. Projections existed between every ordered pair of modules, not only between adjacent ones. All weights were fixed at initialisation.

#### Dynamics

Writing *v*_*a*_(*t*) the membrane potential of neuron *a* at step *t*, the synaptic input was

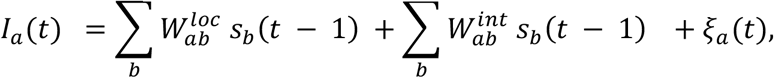

with Poisson drive *ξ*_*a*_(*t*), with a uniform axonal delay *D* = 3 steps (3 ms) on all between-module projections, implemented with a circular spike buffer, and the minimal one-step delay on local recurrence. The membrane followed

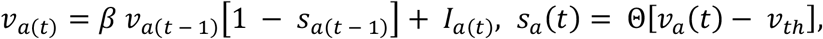

with *β* = 0.9 (*τ*_*m*_ = −*dt* / *ln β* ≈ 9.5 *ms*), *v*_*th*_ = 1.0, reset to zero after a spike, and no refractory period. Potentials and the delay buffer were initialised at zero and each network was simulated for *T* = 10,000 steps (10 s).

#### Drive configurations

The only source of stochastic input was an external Poisson drive *ξ*_*a*_(*t*), implemented as independent Bernoulli spikes of unit amplitude with per-step probability *r* (the base rate). Three spatial configurations were used:

- uniform: every neuron in every module was driven at rate *r*;
- thalamic: only neurons in module 1 were driven at rate *r*, the remaining modules receiving no external input;
- mixed: module 1 was driven at rate *r* and all other modules at *r*/5.

Because the drive is independent across neurons and time, any pairwise or directional structure in the resulting spike trains is produced by the recurrent connectivity alone.

#### Configurations

Networks were generated over the grid shown in Table 1; each combination was instantiated 10 times with independent random connectivity and independent drive, giving an independent ground-truth weight matrix per repeat. The constraint *p*_*fb*_ < *p*_*ff*_ leaves 8 valid pairs across feedforward and feedback strengths.

**Table 1.** Grid of network configurations for the synthetic hierarchy analysis.

| **Parameter** | **Values** |
|---|---|
| Number of modules *B* | 3, 4, 5, 6 |
| Drive configuration | uniform, thalamic, mixed |
| Base rate *r* (Hz) | 1, 2, 3, 4, 5, 10, 20 |
| Feedforward probability *p*<sub>*ff*</sub> | 0.03, 0.04, 0.05 |
| Feedback probability *p*<sub>*fb*</sub> | 0.01, 0.02, 0.03, restricted to *p*<sub>*fb*</sub> < *p*<sub>*ff*</sub> |
| Repeats per configuration | 10 |

### Estimation of module-level connectivity and hierarchy scores

#### STDP coefficients

Because of computational constraints, for each simulated network, the spike train was passed once through the Hebbian attention layer of a SPARKS-fixed model with *τ*_*s*_ = 10 *ms*.

#### Shuffle baseline

STDP coefficients are expected to be non-zero for any pair of active neurons simply because of spurious co-activations. To remove this rate-driven component, the spike train was shuffled by drawing an independent random permutation of the time axis for each neuron, which preserves each neuron’s spike count while removing all cross-neuron timing relations. The coefficient matrix ***A*** was recomputed for *n* = 10 such shuffles and averaged into a baseline ***Ā***, and all analyses used the corrected matrix ***A*** − ***Ā***.

#### Reduction to module-level matrices

The corrected matrix was averaged within each (module, module) block,

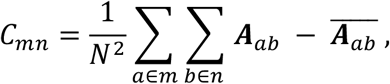

resulting in an *B* × *B* matrix in which *C*_*mn*_ is the mean corrected coefficient for pairs with presynaptic neuron in module *m* and postsynaptic neuron in module *n*. Because the overall scale of the coefficients depends on firing rate and recording length, ***C*** was normalised by the sum of its entries. The same reduction was applied to the ground-truth weight matrix ***W*** = ***W***^loc^ + ***W***^int^**Â**. Entries above the diagonal of normalised matrix Ĉ (respectively Â) are the feedforward coefficients and entries below the diagonal the feedback coefficients.

#### Projection matrix and hierarchy score

Each normalised matrix was anti-symmetrised as

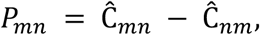

so that *P*_*mn*_ > 0 whenever the influence of module *m* on module *n* exceeds the reciprocal influence. Entries of ***P*** above the diagonal are reported as projection coefficients. The hierarchy score of a module is the sum of its row of the projection matrix,

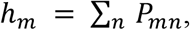

i.e., the net directed influence a module exerts on the rest of the network. Since ***P*** is antisymmetric, the scores sum to zero by construction, and in the ground-truth network, they order the modules along the generative hierarchy. Ground-truth scores were obtained by applying the identical antisymmetrisation and row-sum to Â.

#### Statistics

Inferred and ground-truth quantities were compared per configuration cell. For a given (drive, *B*, base rate), the feedforward, feedback and projection coefficients and the hierarchy scores were concatenated across the 8 (*p*_*fb*_, *p*_*ff*_) pairs and 10 repeats of that cell, and the agreement between inferred and actual values was quantified by the Pearson correlation coefficient and its two-sided *p*-value. The same quantities were additionally pooled across base rates to give one correlation per (*B*, drive) combination. Each repeat contributes *B* hierarchy scores and *B*(*B* − 1)/2 coefficients of each directional type, all from independently drawn networks.

#### Cross-correlogram control

To test whether the hierarchy recovered from the Hebbian coefficients could equally be recovered by an established measure, the same synthetic networks were analysed with jitter-corrected cross-correlograms, following previous work^57^.

For each pair of units,

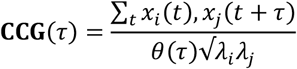

where *θ*(*τ*) = T − |τ| corrects for the shrinking overlap of the sliding window and lambda is a unit’s mean firing rate. From this was subtracted the expected cross-correlogram under 25 ms pattern jitter, which preserves each unit’s spike count within every jitter window and the shape of its rate profile, so that correlations slower than the window cancel and only fast, putatively direct interactions remain.

Following previous work^57^, a pair was counted as connected when the peak of the jitter-corrected cross-correlogram within 10 ms of zero lag exceeded seven times the standard deviation of its flanks, measured at 50-100 ms from zero, and the sign of the peak lag determined which unit led.

#### Cross-correlogram hierarchy

Module-level directionality was summarised by the directionality score introduced previously^57^,

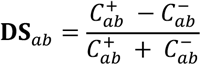

where *C*^+^ counts pairs whose peak lag has a unit in module *a* leading a unit in module *b*, and *C*^−^ counts the reverse. DS is antisymmetric by construction and bounded by 1, so it takes the place of the projection matrix in our analysis. Each module’s hierarchy score is the row mean over its partners, as for the Hebbian coefficients.

#### Recovery of module ranks

For each run, modules were ranked by their inferred and, separately, by their ground-truth hierarchy scores, rank 0 being assigned to the module with the highest score (the most driving module) and rank *B* − 1 to the lowest. A module was counted as correctly placed when its inferred rank equalled its true rank. For each combination, the fraction of correctly placed modules was computed over all modules of all runs of that combination, pooled all possible pairs and repeats. To locate where errors arise in the hierarchy, placement accuracy was also computed separately for each true rank *k* as the fraction of runs in which the module of true rank *k* was assigned inferred rank *k*; because each run contains exactly one module of each true rank, this is a proportion over runs. Rank-wise accuracies were computed per drive and averaged across the three drives with equal weight. For both measures, the accuracy expected from a random ordering of the modules is 1/*B*.

#### Comparison with the cross-correlogram control

Hierarchies inferred from STDP coefficients were compared with those inferred from directed cross-correlograms. Because the modules of one run are not independent (misplacing one module displaces another) comparisons were made per run throughout. The fraction of correctly placed modules was compared for all possible combinations with a two-sided permutation test on the difference in the mean placement fraction per-run, permuting labels within all base rates (10^f^ permutations). Placement accuracy at each true rank, comprising a single binary outcome for every run, was compared across drives with a Cochran–Mantel–Haenszel test by drive and base rate, with continuity correction. Agreement between inferred and ground-truth hierarchy scores was assessed in two ways. The Pearson correlation of scores pooled across the runs of each combination was Fisher-transformed, its standard error estimated by resampling runs with replacement within base rates (2,000 bootstrap resamples), and the two methods compared with a two-sided *z*-test on the difference. The 12 combinations were combined into an overall test by inverse-variance weighting of the *z*-differences.

### Functional Hierarchy of Mouse Visual Cortical Areas

#### Electrophysiological data

We analysed extracellular recordings from the Allen Brain Observatory Visual Coding Neuropixels dataset: 31 sessions in awake, head-fixed mice, recorded simultaneously across six visual cortical areas (VISp, VISal, VISrl, VISpm, VISam, VISl).

Units were included if they passed a signal-to-noise threshold of 1.5 and had at least one spike in every stimulus condition used, the latter so that trial indices stay aligned across the population. This gave 93-386 simultaneously recorded units per session. Because probe placement differs between animals, not every area is present in every session, and we ignored neurons from an area if fewer than 15 were recorded in a session. We used responses to the drifting- and static-grating and flash conditions. Spike trains were binned at *dt* = 1 ms and binarised, and responses were cut into 1132-1360 trials per session.

#### Hierarchy score

Chance-level correction and area-level connectivity were computed following the same protocol as for synthetic data. Area matrices were pooled across sessions by averaging each pair over the sessions in which both areas were present. Each session therefore contributed equally regardless of how many units it happened to record, and pairs never jointly recorded contribute nothing. The number of contributing sessions and the standard error across sessions were retained for every pair. Each session’s matrix was divided by the standard deviation of its own chance-corrected coefficients, taken over all off-diagonal unit pairs in that session; values are therefore expressed in units of the session’s own pairwise coefficient spread. Hierarchy was quantified from the antisymmetric part of the pooled connectivity, following the protocol described above for the synthetic data.

#### Statistics

Confidence intervals were obtained by resampling sessions with replacement (1000 bootstrap replicates), recomputing the pooled connectivity, asymmetries and hierarchy scores within each replicate, and taking the 2.5th and 97.5th percentiles of each score and of each area’s rank.

Sessions are the unit of replication throughout. Trials within a session were used only to estimate that session’s coefficients, never as independent samples, since they share an animal, a probe placement and a unit yield. For individual pairs, the asymmetry was tested against zero with a one-sample t-test across the sessions contributing to that pair.

The recovered ordering was compared against the published anatomical hierarchy of mouse visual cortex (VISp < VISl < VISrl < VISal < VISpm < VISam^57,59^) by Spearman rank correlation between the hierarchy score and the published rank, with a confidence interval from the same session-level bootstrap. This comparison was used only to evaluate the recovered ordering and played no part in producing it.

**Figure S1.**
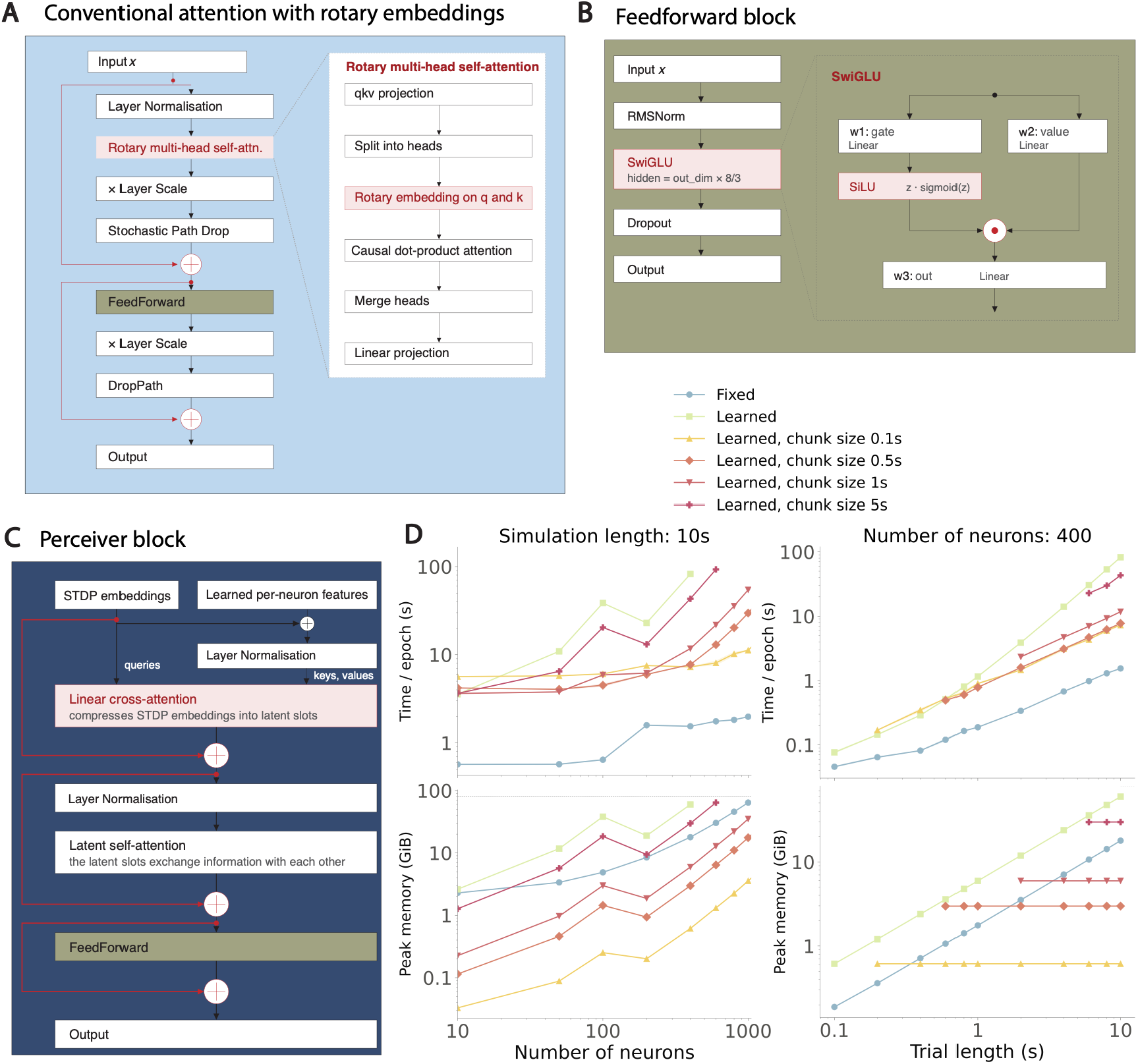
(**A**) Schematic of the conventional dot-product attention block, with layer normalisation^86^, rotary multi-head self-attention^28^, dropout^87^ and dropped paths^72^. qkv: query, key, value. Red: residual paths. (**B**) Schematic of the feedforward network in each attention block, with RMSNorm^88^, SwiGLU^89^ expansion, and dropout. (**C**) Detailed view of the spatial Perceiver-like^73^ block. Red: residual paths. (**D**) Comparison between the fixed and learned configurations training on spike trains simulated with temporal resolution dt = 0.001s on a single NVIDIA A100 GPU. Left: evolution as a function of a number of neurons in the simulation for a fixed simulation length of 10s. Right: evolution as a function of the simulation length for N = 400 neurons. Top: average time per training epoch, calculated over 20 epochs. Bottom: Average peak memory consumption, calculated over 20 epochs. Shaded areas represent standard deviation over 20 epochs.

**Figure S2.**
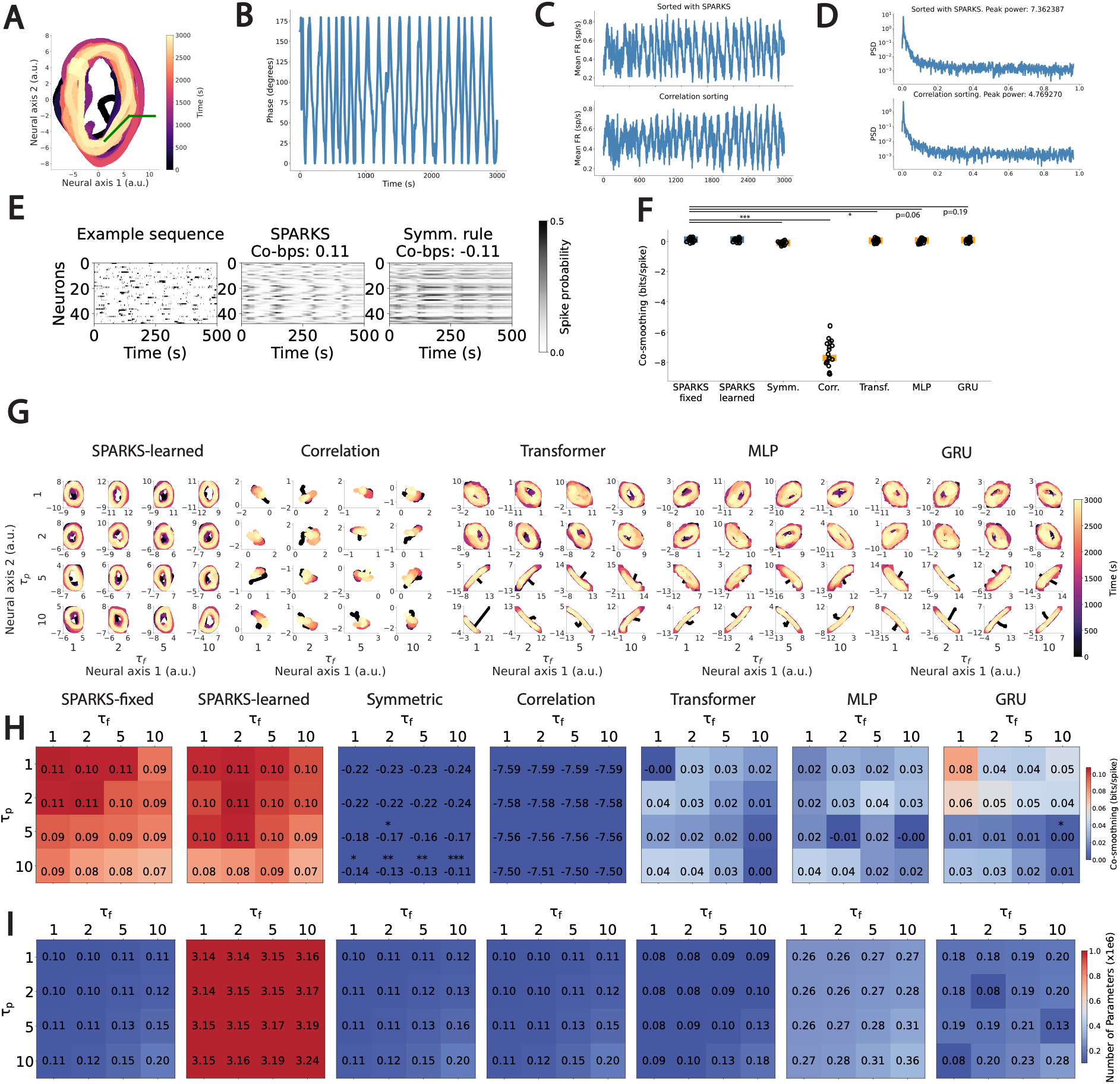
(**A**) Illustration of the angle used to recover the phase of the oscillations from the latent embeddings. (**B**) Phase signal recovered from the latent embeddings obtained with SPARKS-fixed. (**C**) Oscillatory signals recovered by population decoding of the spike trains after sorting neurons with different methods. (**D**) PSDs corresponding to signals in (**C**). (**E**) Raster plot for left-out neurons (left) and corresponding predicted spike probabilities after training with SPARKS-fixed (centre), and the benchmark symmetric rule (right). (**F**) Best co-smoothing scores obtained for SPARKS and control models for which we replaced the Hebbian attention block with a different deep learning module. Symm: symmetric Hebbian attention, MLP: multilayer perceptron, GRU: gated recurrent unit, Corr: correlation-based module (Methods), Transf.: conventional Transformer block ^27^. (**G**) Latent embeddings obtained with SPARKS and control modules across different values of τ_f_ and τ_p_. (**H**) Evolution of the co-smoothing loss for different values of τ_f_ and τ_p_ for SPARKS and all control. Stars indicate statistical difference from the scores obtained for τ_p_ = τ_f_ = 1 timestep, measured by a two-sided Mann-Whitney U test. *: p < 0.05, **:p < 0.01, ***:p < 0.001. (**I**) Number of parameters for SPARKS and control modules across different values of τ_f_ and τ_p_.

**Figure S3.**
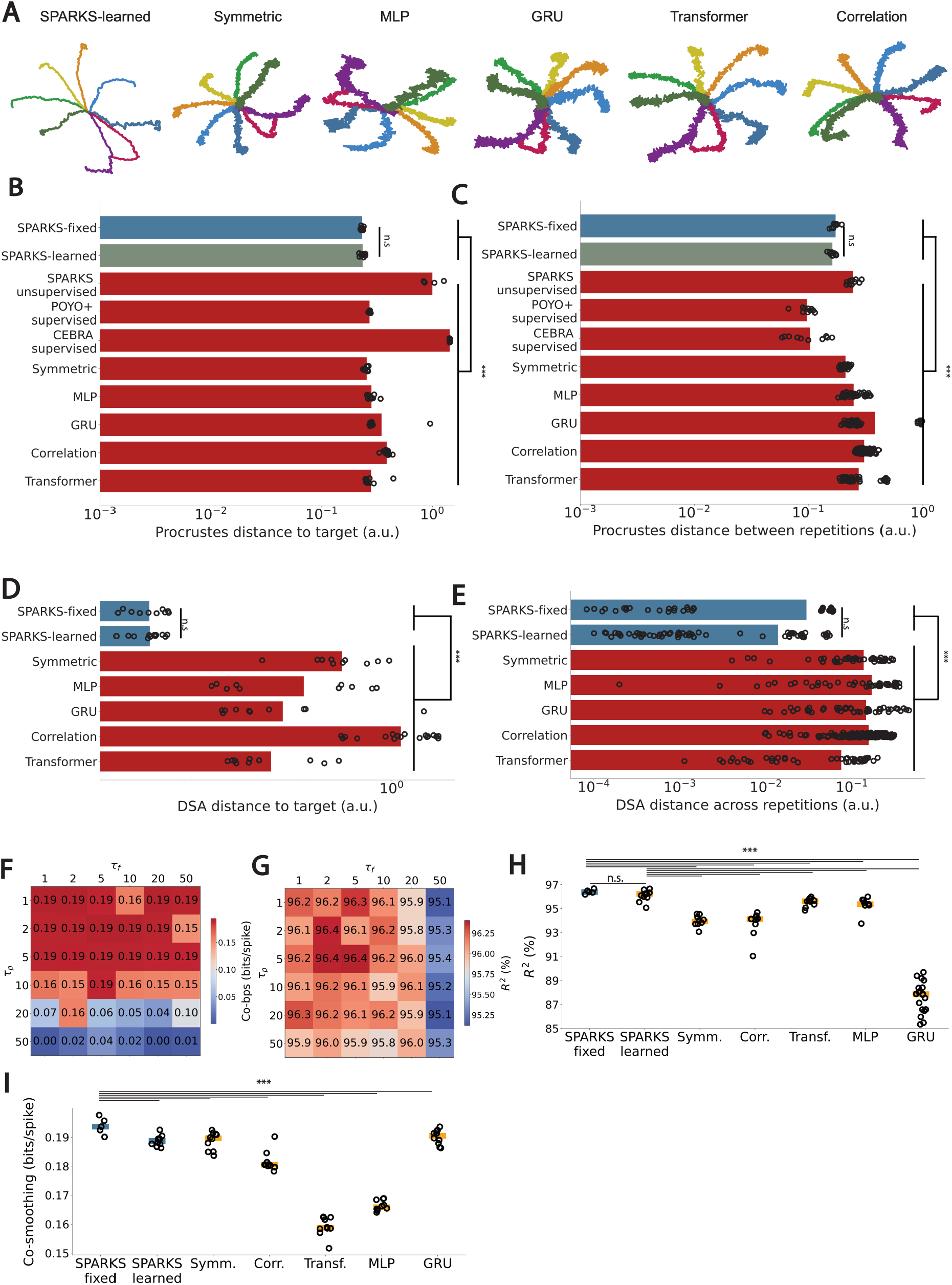
(**A**) Example latent embeddings for models trained via supervised learning. (**B**) Comparison of the Procrustes distances between latent embeddings and the task parameter. Each value represents the mean over 10 individual runs, circles represent individual runs. (**C**) Comparison of the Procrustes distances between the latent embeddings across 10 repetitions after training with random weights initialisation for each model type. Each value represents the mean value, circles represent individual runs. (**D**) Comparison of the DSA distances between latent embeddings and the task parameter. Each value represents the mean over 10 individual runs, circles represent individual runs. (**E**) Comparison of the DSA distances between the latent embeddings across 10 repetitions after training with random weights initialisation for each model type. Each value represents the mean value, circles represent individual runs. (**F**), (**G**) Scores for different values of τ_p_ and τ_f_ for SPARKS-fixed for co-smoothing and supervised prediction of the animal’s hand position. (**H**) Comparison of R^2^ scores for prediction of hand position. Horizontal lines represent the median over 20 runs, and each dot an individual run. (**I**) Comparison of co-smoothing scores. Horizontal lines represent the median over 5 runs, and each dot an individual run. *: p < 0.05, **:p < 0.01, ***:p < 0.001, n.s.: not significant, two-sided Mann-Whitney U-test.

**Figure S4.**
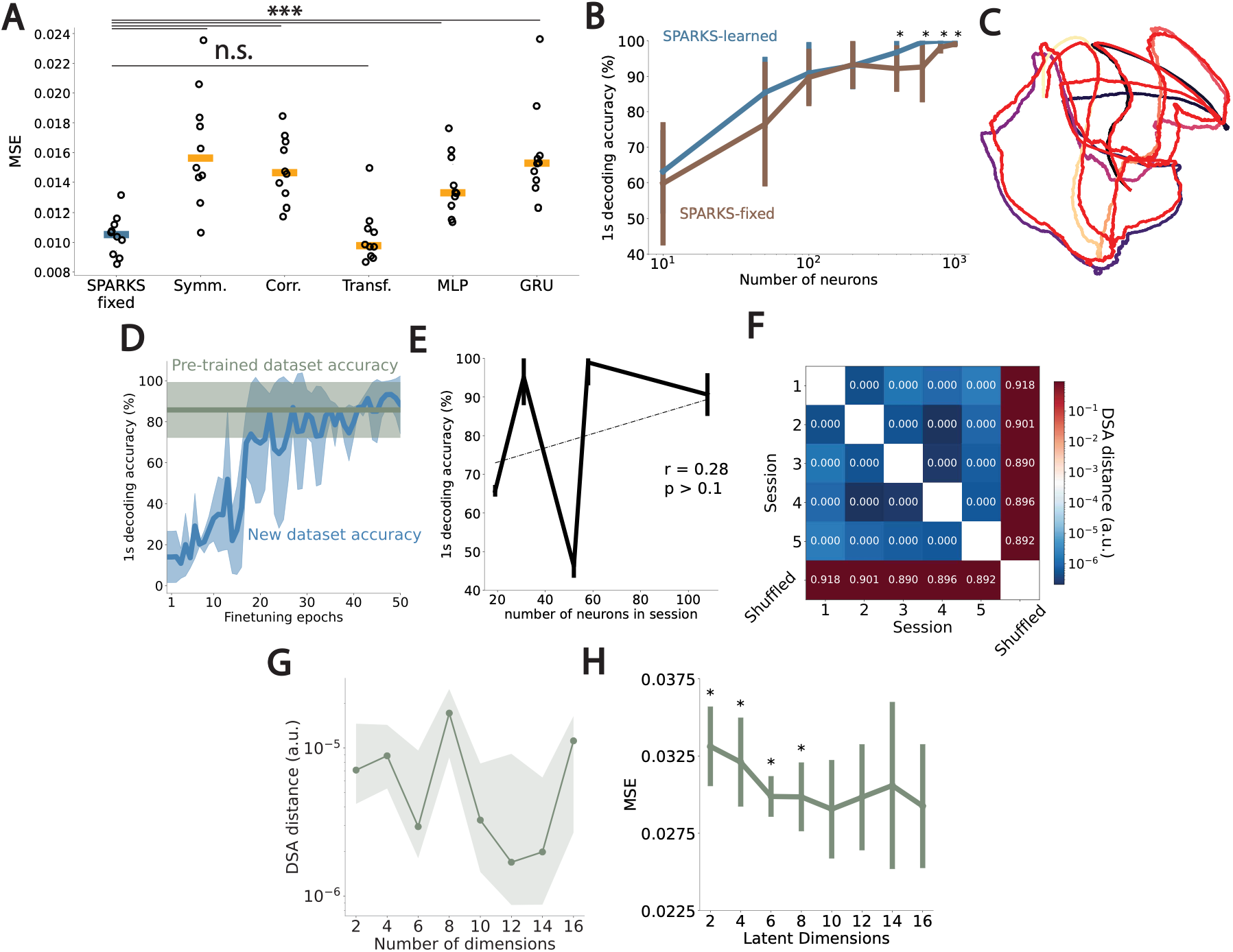
(**A**) Comparison of MSE scores for movie reconstruction. Horizontal lines represent the median over 10 runs, and each dot an individual run. (**B**) 1s decoding accuracy as a function of number of randomly sampled neurons for SPARKS-learned and SPARKS-fixed. Error bars represent standard deviation across 10 repetitions (**C**) Latent embeddings averaged across examples from blocks 1 and 2 for a SPARKS-fixed model trained on block 1 only. (**D**) Frame decoding accuracy as a function of finetuning epochs for SPARKS-fixed. Green trace corresponds to the pre-trained dataset. Blue trace shows accuracy for a newly introduced dataset to an existing model. Shaded areas represent standard deviation over 5 pre-trained models. (**E**) 1s decoding accuracy as a function of number of neurons in a session for SPARKS-fixed. Error bars represent standard deviation across 5 repetitions. r: Pearson correlation coefficient. p: p-value. (**F**) Pairwise DSA distances between latent embeddings obtained for all sessions for unsupervised learning with SPARKS-fixed, average across 5 repetitions. Shuffled: latent representations from session 0, randomly shuffled across the time dimension. (**G**) Average pairwise DSA distances between latent embeddings of five different sessions obtained from SPARKS-fixed as a function of the latent dimensionality. Shaded areas represent interquartile range across 5 repetitions and sessions. (**H**) Average reconstruction MSE across sessions from SPARKS-fixed trained via unsupervised learning as a function of the latent dimensionality. Error bars represent standard deviation across repetitions and sessions. Stars represent statistical difference with the minimum obtained at 10 latent dimensions. *: p < 0.05, **: p < 0.01, *** : p < 0.001.

**Figure S5.**
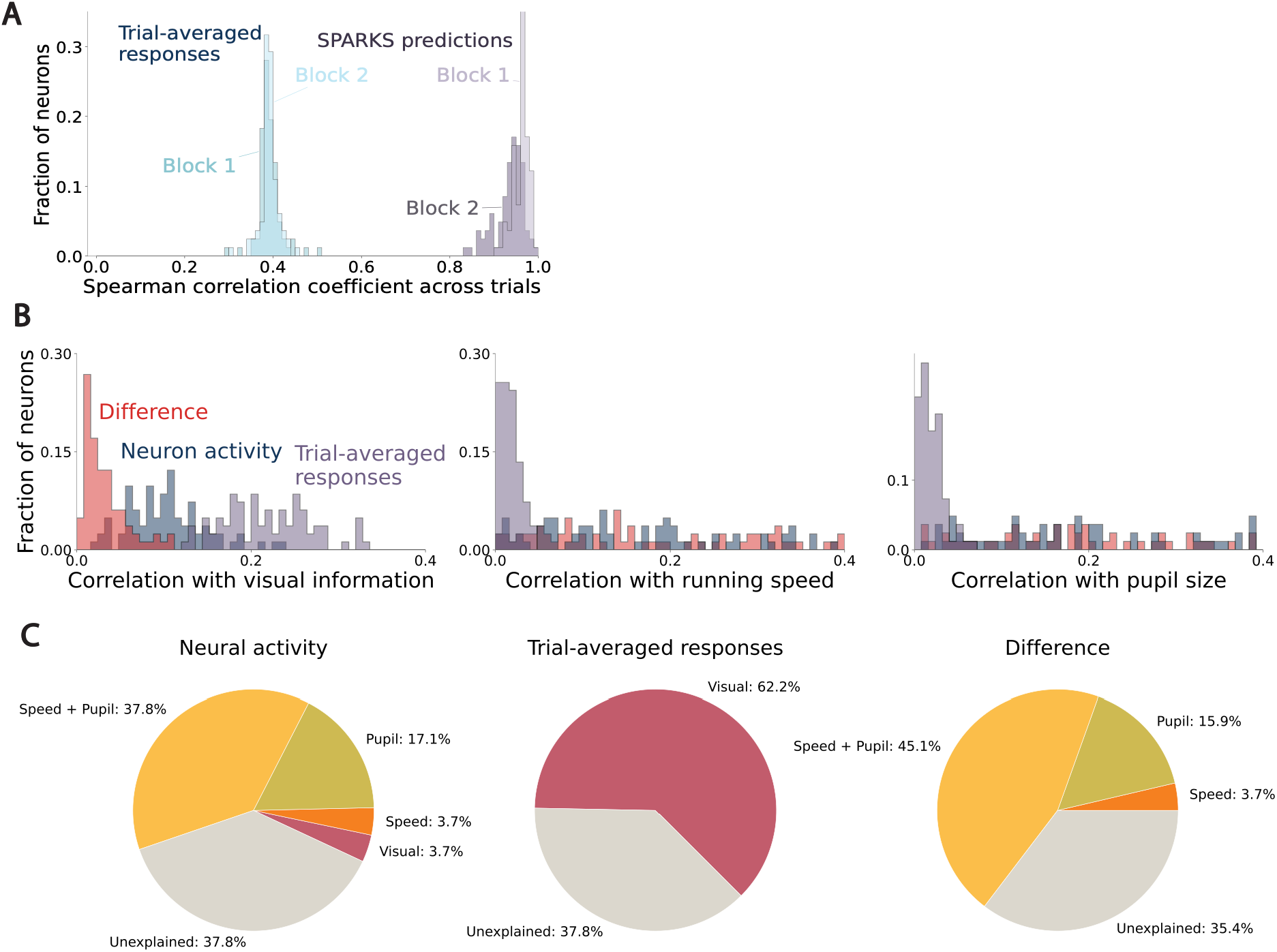
(**A**) Spearman correlation of trial-averaged responses and SPARKS-learned predictions to each neurons’ responses for block 1 and block 2. (**B**) Distribution of correlation coefficients for raw neuronal activity (blue), trial-averaged responses (purple), and for the difference between the two (pink) with pixels brightness (left), running speed (centre), and pupil size (right). (**C**) Pie charts representing the fractions of neuron correlated to measured parameters, based on their activity (left), Trial-averaged responses (centre), and the difference between the two (right).

**Figure S6.**
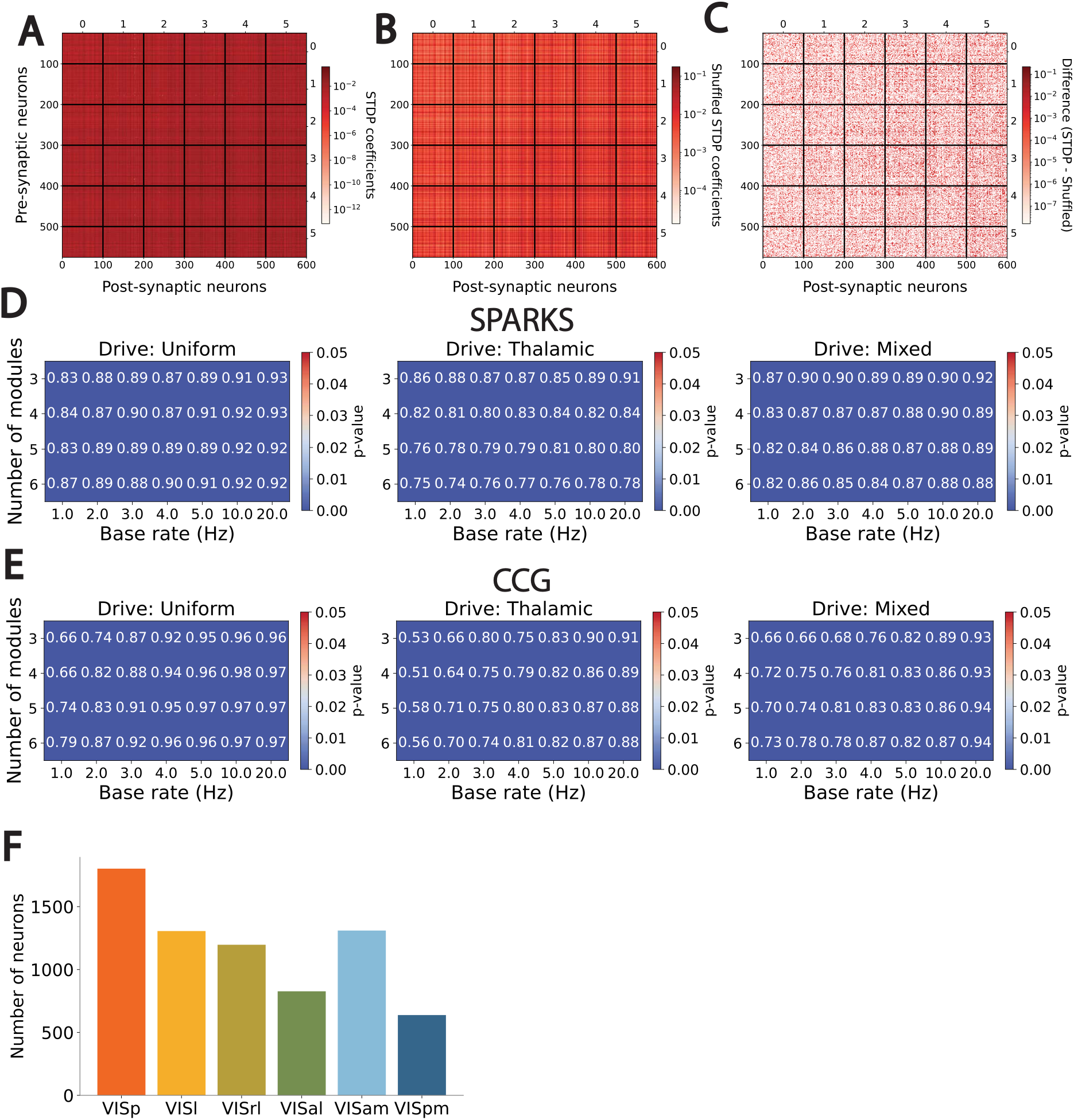
(**A**) STDP coefficients obtained from the Hebbian attention layer in SPARKS-fixed at the end of an example for a representative network comprising six modules. Vertical and horizontal lines represent separation between modules. (**B**) STDP coefficients obtained from the Hebbian attention layer in SPARKS-fixed at the end of a shuffled example for a representative network comprising six modules. Sequences were shuffled for each neuron independently. Vertical and horizontal lines represent separation between modules (**C**) Difference between STDP coefficients obtained from the Hebbian attention layer in SPARKS-fixed for the actual and shuffled example from a representative network comprising six modules. Sequences were shuffled for each neuron independently. Vertical and horizontal lines represent separation between modules. (**D**) Correlations between the actual hierarchy and the hierarchy predicted with SPARKS-fixed. (**E**) Correlations between the actual hierarchy and the hierarchy obtained based on CCGs^57^. (**F**) Total number of neurons for each HVA in the Allen visual coding dataset.

**Table S1.**
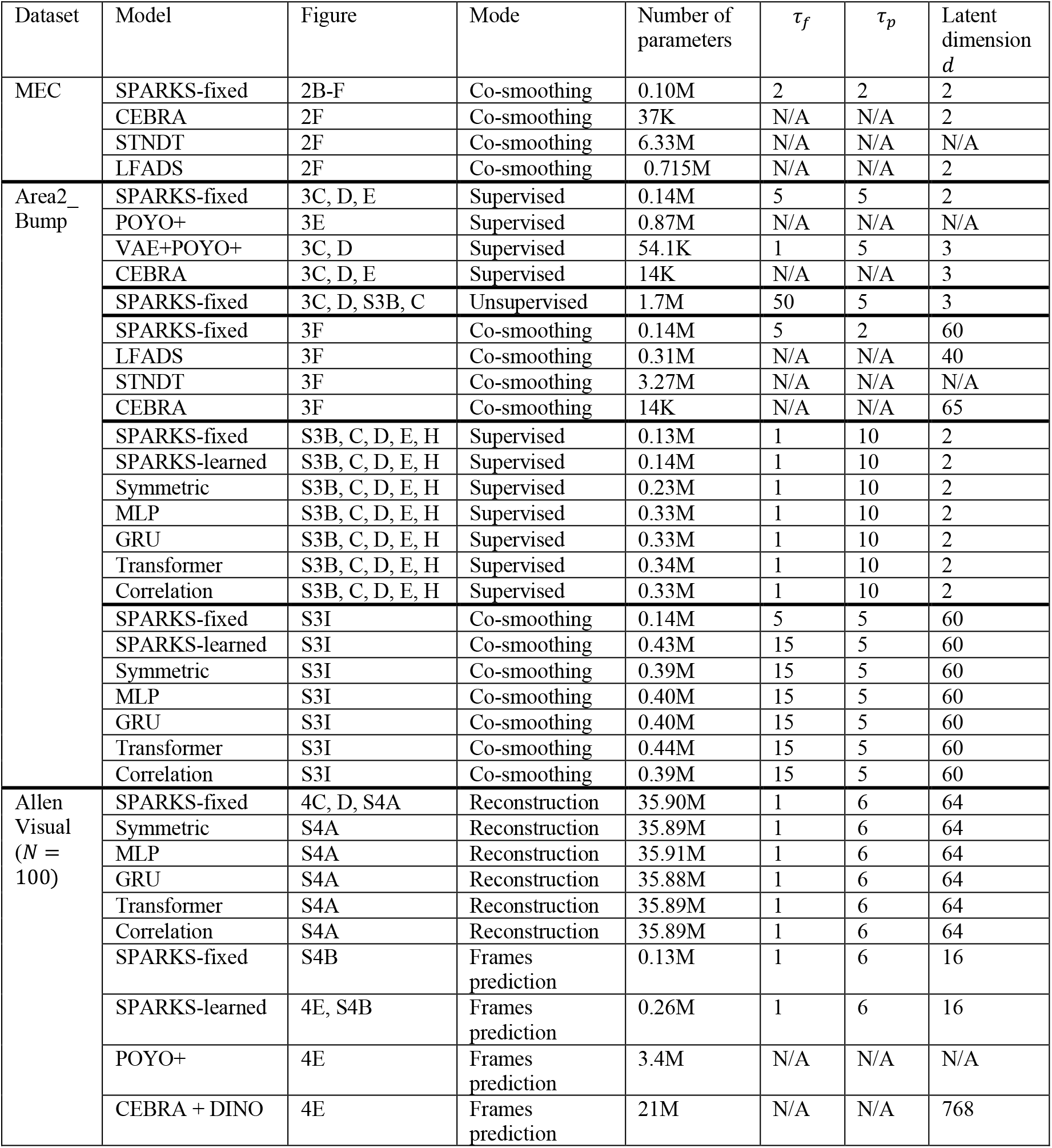
Summary of parameters used for comparisons with competing models.

| Dataset | Model | Figure | Mode | Number of parameters | τ<sub>f</sub> | τ<sub>p</sub> | Latent dimension *d* |
|---|---|---|---|---|---|---|---|
| MEC | SPARKS-fixed | 2B-F | Co-smoothing | 0.10M | 2 | 2 | 2 |
|  | CEBRA | 2F | Co-smoothing | 37K | N/A | N/A | 2 |
|  | STNDT | 2F | Co-smoothing | 6.33M | N/A | N/A | N/A |
|  | LFADS | 2F | Co-smoothing | 0.715M | N/A | N/A | 2 |
| Area2_Bump | SPARKS-fixed | 3C, D, E | Supervised | 0.14M | 5 | 5 | 2 |
|  | POYO+ | 3E | Supervised | 0.87M | N/A | N/A | N/A |
|  | VAE+POYO+ | 3C, D | Supervised | 54.1K | 1 | 5 | 3 |
|  | CEBRA | 3C, D, E | Supervised | 14K | N/A | N/A | 3 |
|  | SPARKS-fixed | 3C, D, S3B, C | Unsupervised | 1.7M | 50 | 5 | 3 |
|  | SPARKS-fixed | 3F | Co-smoothing | 0.14M | 5 | 2 | 60 |
|  | LFADS | 3F | Co-smoothing | 0.31M | N/A | N/A | 40 |
|  | STNDT | 3F | Co-smoothing | 3.27M | N/A | N/A | N/A |
|  | CEBRA | 3F | Co-smoothing | 14K | N/A | N/A | 65 |
|  | SPARKS-fixed | S3B, C, D, E, H | Supervised | 0.13M | 1 | 10 | 2 |
|  | SPARKS-learned | S3B, C, D, E, H | Supervised | 0.14M | 1 | 10 | 2 |
|  | Symmetric | S3B, C, D, E, H | Supervised | 0.23M | 1 | 10 | 2 |
|  | MLP | S3B, C, D, E, H | Supervised | 0.33M | 1 | 10 | 2 |
|  | GRU | S3B, C, D, E, H | Supervised | 0.33M | 1 | 10 | 2 |
|  | Transformer | S3B, C, D, E, H | Supervised | 0.34M | 1 | 10 | 2 |
|  | Correlation | S3B, C, D, E, H | Supervised | 0.33M | 1 | 10 | 2 |
|  | SPARKS-fixed | S3I | Co-smoothing | 0.14M | 5 | 5 | 60 |
|  | SPARKS-learned | S3I | Co-smoothing | 0.43M | 15 | 5 | 60 |
|  | Symmetric | S3I | Co-smoothing | 0.39M | 15 | 5 | 60 |
|  | MLP | S3I | Co-smoothing | 0.40M | 15 | 5 | 60 |
|  | GRU | S3I | Co-smoothing | 0.40M | 15 | 5 | 60 |
|  | Transformer | S3I | Co-smoothing | 0.44M | 15 | 5 | 60 |
|  | Correlation | S3I | Co-smoothing | 0.39M | 15 | 5 | 60 |
| Allen Visual (*N* = 100) | SPARKS-fixed | 4C, D, S4A | Reconstruction | 35.90M | 1 | 6 | 64 |
|  | Symmetric | S4A | Reconstruction | 35.89M | 1 | 6 | 64 |
|  | MLP | S4A | Reconstruction | 35.91M | 1 | 6 | 64 |
|  | GRU | S4A | Reconstruction | 35.88M | 1 | 6 | 64 |
|  | Transformer | S4A | Reconstruction | 35.89M | 1 | 6 | 64 |
|  | Correlation | S4A | Reconstruction | 35.89M | 1 | 6 | 64 |
|  | SPARKS-fixed | S4B | Frames prediction | 0.13M | 1 | 6 | 16 |
|  | SPARKS-learned | 4E, S4B | Frames prediction | 0.26M | 1 | 6 | 16 |
|  | POYO+ | 4E | Frames prediction | 3.4M | N/A | N/A | N/A |
|  | CEBRA + DINO | 4E | Frames prediction | 21M | N/A | N/A | 768 |

**Table S2.** Overview of correlations to actual hierarchy scores. Statistical differences were assessed based on a two-sided *z*-test (Methods).

| **Modules** | **Drive** | **SPARKS** | **CCG** | ***p*** |
|---|---|---|---|---|
| **3** | Uniform | **0.88** | 0.74 | 5.3E-54 |
| **3** | Thalamic | **0.87** | 0.77 | 5.4E-23 |
| **3** | Mixed | **0.90** | 0.61 | 1.4E-212 |
| **4** | Uniform | **0.89** | 0.77 | 6.1E-61 |
| **4** | Thalamic | **0.82** | 0.75 | 1.1E-14 |
| **4** | Mixed | **0.87** | 0.63 | 3.5E-223 |
| **5** | Uniform | **0.89** | 0.80 | 2.7E-46 |
| **5** | Thalamic | **0.79** | 0.77 | 0.04 |
| **5** | Mixed | **0.86** | 0.64 | 7.9E-193 |
| **6** | Uniform | **0.89** | 0.79 | 8.9E-68 |
| **6** | Thalamic | **0.76** | **0.76** | 0.52 |
| **6** | Mixed | **0.85** | 0.66 | 5.59E-157 |

**Table S3.** Fraction of correctly recovered ranks in the hierarchy. Statistical differences were assessed with a two-sided permutation test (Methods).

| **Modules** | **Drive** | **SPARKS** | **CCG** | **Difference** | ***p*** |
|---|---|---|---|---|---|
| **3** | Uniform | **0.88** | **0.87** | 0.01 | 0.54 |
|  | Thalamic | **0.81** | 0.76 | 0.05 | 0.01 |
|  | Mixed | **0.86** | 0.76 | 0.1 | 9.9E-06 |
| **4** | Uniform | 0.78 | **0.83** | -0.05 | 0.002 |
|  | Thalamic | **0.68** | **0.65** | 0.03 | 0.08 |
|  | Mixed | **0.78** | 0.72 | 0.06 | 0.0003 |
| **5** | Uniform | 0.71 | **0.79** | -0.08 | 9.9E-06 |
|  | Thalamic | **0.62** | **0.60** | 0.02 | 0.22 |
|  | Mixed | **0.7** | 0.63 | 0.07 | 3.9E-05 |
| **6** | Uniform | 0.63 | **0.72** | -0.09 | 9.9E-06 |
|  | Thalamic | **0.56** | 0.51 | 0.05 | 0.0001 |
|  | Mixed | **0.67** | 0.61 | 0.06 | 6.99E-05 |

**Table S4.** Fraction of correctly placed modules per actual rank in the hierarchy. Statistical differences were assessed with a Cochran–Mantel–Haenszel test, with continuity correction (Methods).

| **Modules** | **Rank** | **SPARKS** | **CCG** | **Difference** | ***p*** |
|---|---|---|---|---|---|
| **3** | 0 | **0.96** | 0.87 | 0.09 | 6.08E-18 |
|  | 1 | **0.78** | 0.72 | 0.06 | 0.0001 |
|  | 2 | **0.82** | **0.80** | 0.02 | 0.12 |
| **4** | 0 | **0.92** | 0.85 | 0.07 | 1.98E-11 |
|  | 1 | **0.68** | **0.66** | 0.02 | 0.22 |
|  | 2 | 0.62 | **0.65** | -0.03 | 0.029 |
|  | 3 | **0.77** | **0.76** | 0.001 | 1.0 |
| **5** | 0 | **0.87** | 0.81 | 0.06 | 5.48E-07 |
|  | 1 | **0.62** | **0.62** | -0.001 | 0.95 |
|  | 2 | 0.53 | **0.58** | -0.05 | 0.008 |
|  | 3 | **0.59** | **0.60** | -0.01 | 0.63 |
|  | 4 | **0.76** | **0.74** | 0.02 | 0.058 |
| **6** | 0 | **0.86** | 0.79 | 0.07 | 3.10E-11 |
|  | 1 | **0.54** | **0.55** | -0.01 | 0.86 |
|  | 2 | 0.46 | **0.51** | -0.05 | 0.031 |
|  | 3 | **0.53** | **0.53** | -0.00 | 0.82 |
|  | 4 | **0.59** | **0.58** | 0.01 | 0.22 |
|  | 5 | **0.74** | 0.72 | 0.02 | 0.19 |

**Movie S1**.

Left: Averaged activity of 200 neurons in response to a naturalistic movie. Right, top: naturalistic movie and bottom: corresponding reconstruction using 200 neurons recorded from mouse primary visual cortex.

